# A Two-Player Iterated Survival Game

**DOI:** 10.1101/465294

**Authors:** John Wakeley, Martin Nowak

**Affiliations:** Department of Organismic and Evolutionary Biology, Harvard University, Cambridge, MA, 02138, USA; Program for Evolutionary Dynamics, Harvard University, Cambridge, MA 02138, USA; Department of Mathematics, Harvard University, Cambridge, MA 02138, USA

**Keywords:** Prisoner’s Dilemma, Survival Game, Iterated Game, Replicator Equation, Moran model

## Abstract

We describe an iterated game between two players, in which the payoff is to survive a number of steps. Expected payoffs are probabilities of survival. A key feature of the game is that individuals have to survive on their own if their partner dies. We consider individuals with hardwired, unconditional behaviors or strategies. When both players are present, each step is a symmetric two-player game. The overall survival of the two individuals forms a Markov chain. As the number of iterations tends to infinity, all probabilities of survival decrease to zero. We obtain general, analytical results for *n*-step payoffs and use these to describe how the game changes as *n* increases. In order to predict changes in the frequency of a cooperative strategy over time, we embed the survival game in three different models of a large, well-mixed population. Two of these models are deterministic and one is stochastic. Offspring receive their parent’s type without modification and fitnesses are determined by the game. Increasing the number of iterations changes the prospects for cooperation. All models become neutral in the limit (*n* → ∞). Further, if pairs of cooperative individuals survive together with high probability, specifically higher than for any other pair and for either type when it is alone, then cooperation becomes favored if the number of iterations is large enough. This holds regardless of the structure of pairwise interactions in a single step. Even if the single-step interaction is a Prisoner’s Dilemma, the cooperative type becomes favored. Enhanced survival is crucial in these iterated evolutionary games: if players in pairs start the game with a fitness deficit relative to lone individuals, the prospects for cooperation can become even worse than in the case of a single-step game.

## 1. Introduction

More than a century ago, Kropotkin (1902) argued that what he called mutual aid should be ranked among the main factors of evolution, an even more important driver of evolution than within-species competition. The idea of mutual aid is similar to that of mutualism: partnerships may be beneficial to both partners and thus unconditionally favored. Kropotkin had been deeply impressed by the abilities of animals to endure harsh winters and perilous migrations in northern Eurasia by living together and supporting each other in situations where a lone animal had little chance of surviving. Thus, he inferred a connection between cooperative behaviors and survival under adverse conditions. But Kropotkin did not spell out the ways in which adversity might favor cooperative behaviors, nor did he consider that cooperative behaviors could be disadvantageous. He took the very fact that animals seem to do better in groups than alone as evidence for mutual aid, apparently unaware that any interaction presents the opportunity, or at least the temptation, to be the one who comes out ahead even if it is at the expense of other individuals.

Indeed, game theory and evolutionary theory have uncovered numerous situations in which cooperative or otherwise helping behaviors are detrimental to the individual or selectively disadvantageous (Hofbauer and Sigmund, 1998; Mesterton-Gibbons, 2000; Cressman, 2005; Schecter and Gintis, 2016). It has been shown using a variety of models that such behaviors can be disfavored even if there are obvious advantages to mutual cooperation. For example, in the classic two-player game called the Prisoner’s Dilemma (Tucker, 1950; Rapoport and Chammah, 1965; Axelrod, 1984), cooperation results in a “reward payoff” if one’s partner also cooperates but a “sucker’s payoff” if one’s partner defects. Defection yields a “temptation payoff” if one’s partner cooperates but a “punishment payoff” if one’s partner also defects. The reward is of course better than the punishment, but it is further assumed that the temptation payoff is greater than the reward, and the sucker’s payoff is even worse than the punishment. This leads to a paradox: an individual who defects always receives a higher payoff but it is clearly illogical for everyone to defect.

The Prisoner’s Dilemma, in which the payoff for defection is higher than for cooperation regardless of one’s partner behavior, represents one of four possible types of two-player games. It is the polar opposite of what Kropotkin (1902) imagined for mutual aid, that instead it would cooperation that would always yield the higher payoff. Between these extremes lie two other kinds of games. In the Stag Hunt (Skyrms, 2004) and related games, the payoff to an individual is higher if it matches the partner’s behavior regardless of what that behavior is. Cooperators in the Stag Hunt go for the big game, a stag, which two such individuals can catch but one alone cannot. Non-cooperators opt out of the big-game hunt and accept a middling payoff, a hare, which a single individual can catch. In this case, cooperation yields the higher payoff only when one’s partner also cooperates. The Hawk-Dove game (Maynard Smith and Price, 1973; Maynard Smith, 1978) represents the fourth kind, in which having a different behavior than one’s partner produces a higher payoff. Doves cooperate by sharing resources but retreat when challenged. Hawks don’t cooperate. They are ready to fight to avoid leaving an interaction empty-handed. A Hawk gets the entire resource when facing a Dove but suffers badly when facing another Hawk. Hawk and Dove, of course, refer to stereotypical personalities not animals. In this last case, cooperation yields the higher payoff only when one’s partner does not cooperate. Besides Hawk-Dove, other well-studied games of this type are the game of chicken and the snowdrift game (Doebeli and Hauert, 2005; Nowak, 2006a).

We introduce a two-player survival game which can fall into any of these four classes. It may also change, for example from a Hawk-Dove game into a Prisoner’s Dilemma, depending on one key feature, the length of the game. In this game, an initially sampled pair of individuals confronts a hazardous situation which is repeated *n* times. With reference to Kropotkin (1902), we might imagine that each day of a long journey presents a similar set of challenges which threaten survival. They might, for example, be trying to survive a number of very cold nights or attempting to defend themselves repeatedly against a predator. How they fare in each step depends on their behavior and their partner’s behavior. The payoff for an individual is all or nothing: either survive to the end of the game or not. Crucially, an individual must face the perilous situation alone for the remainder of the game if its partner dies. Survival payoffs are thus meted out at each step, and this will be important for determining total payoffs in the game. Denoting survival as 1 and death as 0, the expected total payoff to an individual in a given situation (i.e. with a specified partner initially) is its probability of surviving to the end of the game. The *n*-step survival game is a symmetric two-player game with total payoffs, *a*(*n*), *d*(*n*), *c*(*n*) and *d*(*n*) as Table 1. Using Table 1 to represent an *n*-step game that is a Prisoner’s Dilemma, with *A* for cooperation and *B* for defection, the reward would be *a*(*n*), the sucker’s payoff *b*(*n*), the temptation payoff *c*(*n*) and the punishment *d*(*n*).

**Table 1:** The payoff, *a*(*n*), *b*(*n*), *c*(*n*) or *d*(*n*), that an individual receives in a symmetric two-player game depends both on the individual and on the individual’s partner. In a survival game, these payoffs are probabilities of survival. *A* and *B* denote two possible strategies or types of individuals (e.g. Dove and Hawk). Payoffs are simultaneously awarded to both players, i.e. each is considered the Individual (and the other the Partner) and awarded a payoff according to the table.

|  |  | Partner |  |
| --- | --- | --- | --- |
| | | $A$ | $B$ |
| Individual | $A$ | $a(n)$ | $b(n)$ |
| | $B$ | $c(n)$ | $d(n)$ |

More generally, depending on *a*(*n*), *b*(*n*), *c*(*n*) and *d*(*n*), any *n*-step survival game falls into one of the four classes described above. Ignoring the detail that some payoffs might be identical, these are defined as follows. If *a*(*n*) < *c*(*n*) and *b*(*n*) < *d*(*n*), cooperation has the lower payoff regardless of the partner. Then the game falls into the class exemplified by the Prisoner’s Dilemma, which we will call PD for short. If *a*(*n*) > *c*(*n*) and *b*(*n*) < *d*(*n*), cooperation has the higher payoff only when one’s partner cooperates. This is the Stag-Hunt class of games, or SH for short. If *a*(*n*) < *c*(*n*) and *b*(*n*) > *d*(*n*), cooperation has the higher payoff only when one’s partner doesn’t cooperate. This is the Hawk-Dove class, or HD for short. Finally, if *a*(*n*) > *c*(*n*) and *b*(*n*) > *d*(*n*), cooperation has the higher payoff regardless of the partner. We follow De Jaegher and Hoyer (2016) and call this a Harmony Game, or HG for short.

We refer to the *n*-step survival game as a repeated or iterated game because the payoff structure in each of the *n* steps is identical and equivalent to the single-step version of the game. The notion of repeated or iterated games is generally predicated on the idea that individuals can act differently in different steps and can react to their partner’s behavior. When this is true, the relatively simple conclusions about which strategy may be favorable based on Table 1 do not necessarily hold. For example, if two individuals play an iterated Prisoner’s Dilemma under these conditions, then reactive strategies like Tit-for-Tat (Axelrod, 1984) or win-stay, lose-shift (Nowak and Sigmund, 1993) can be favored over the single-iteration strategy of always-defect. Such repeated games allow for the phenomena of direct and indirect reciprocity (Trivers, 1971; Nowak and Sigmund, 1998) and trigger strategies which punish non-cooperation (Osborne and Rubinstein, 1994). These are examples of a general phenomenon of behavioral responsiveness (Van Cleve and Akąy, 2014) which is one of a small number of mechanisms known to promote the evolution of costly cooperation (reviewed in Nowak, 2006b; Van Cleve and Akąy, 2014).

We do not consider reactive strategies or behavioral responsiveness in this work. Individuals have one of two possible simple strategies: *A* or *B* as in Table 1. When both individuals are present, which is always the case initially, their survival probabilities in each iteration are given by the single-step version of Table 1, i.e. with *a*(1), *b*(1), *c*(1) and *d*(1). We will refer to these single-step survival probabilities as *a*, *b*, *c* and *d*. The *n*-step survival probabilities, *a*(*n*), *b*(*n*), *c*(*n*) and *d*(*n*), will depend on these as well as on the probabilities of survival when an individual is alone. Although the game always begins with two individuals, if one dies the other must continue. In the remaining steps, a loner plays a game against Nature. Specifically, a loner survives a single step with probability *a*_0_ if it has strategy *A* and probability *d*_0_ if it has strategy *B*. We are especially interested in cases in which the single-step, two-player game defined by *a*, *b*, *c* and d is of a different type than the *n*-step game defined by *a*(*n*), *b*(*n*), *c*(*n*) and *d*(*n*).

Several previous works have considered games in which the random survival of individuals is an important factor and environmental conditions may be harsh. Eshel and Weinshall (1988) introduced a model in which payoffs are probabilities of survival. Payoffs were drawn randomly from a distribution, and the game was repeated with a fixed probability. As in the model we propose, Eshel and Weinshall (1988) allowed that the game may continue even if one individual dies. They considered optimal strategy choice by indivdiuals under the assumption that individuals have perfect knowledge of the game’s structure, including the distribution of the payoffs. Eshel and Shaked (2001) described a similar survival game but included a general probability that both members of a pair survive, thus allowing for arbitrary synergistic effects of partnership (Hauert et al., 2006; Kun et al., 2006) within each step of the game. Eshel and Weinshall (1988) had assumed that pairwise survival probabilities were the products of two individual survival probabilities.

Garay (2009) combined the idea of a survival game with the “selfish herd” theory of Hamilton (1971) to produce a model of cooperation between pairs of individuals in defense against repeated attacks by a predator. Individuals were of two possible types, characterized by a probability of helping their partner in defense against attack. Garay (2009) derived the total survival probabilities of individuals in different kinds of partnerships given a fixed number of attacks, and used these to describe evolutionarily stable strategies (Maynard Smith and Price, 1973) in an infinite population. Using the example contrast of one attack versus eight attacks, this revealed cases in which helping could not invade a completely selfish population given a one-attack game but could invade if the number of attacks was larger (Garay, 2009).

Kropotkin’s notion that high levels of adversity could favor cooperation has also been studied using simulations of structured populations. Harms (2001) found that in a Prisoner’s Dilemma cooperators could gain an advantage at the inhospitable margins of a population by colonizing patches cleared by the local extinction of defectors. More recent simulations of another model by Smaldino et al. (2013) also found cases in which cooperation can be favored despite the Prisoner’s Dilemma. Specifically, if there is a high cost of living for all individuals which, if unchecked, would lead to the extinction of the population and if occasional interactions with cooperators are required for survival, then cooperation can be favored. As in Harms (2001), the success of cooperators in this case relied on their ability to form clusters (Smaldino et al., 2013).

De Jaegher and Hoyer (2016) considered two game-theoretic models of the behavior and ecology of a pair of individuals, in which higher levels of adversity can change the type of the game. Their Model 1 includes a degree of complementarity which rescales payoffs for mixed-strategy pairs compared to same-strategy pairs, and which could represent an environmental challenge for mixed pairs. Their Model 2 includes a number of attacks by a predator on individuals protecting a common resource. Cooperators are immune to attacks while defectors can sustain at most one attack before the common resource is lost. As in Garay (2009), the number of attacks indicates the level of environmental challenge. Cooperation may be disfavored when the number of attacks is small and yet become favored when the number of attacks is large. Depending on the other parameters in the model, however, a range of other switches between types of games may occur as the number of attacks increases. De Jaegher (2017) extended these results to multi-player games.

Our concerns here are similar to those of Garay (2009) and De Jaegher and Hoyer (2016). We study how the structure of the two-player iterated survival game changes as a function of its parameters, in particular the number of steps *n*. We compare the conclusions for single-step games with those for *n*-step games. Because the payoffs which are probabilities of survival in this game decrease as *n* increases, *n* is a measure of adversity. We present general results for any level of adversity, then focus on the possible structures of large-*n* games. We find conditions under which any type of single-step game, be it a Prisoner’s Dilemma, a Stag Hunt or a Hawk-Dove game, will become a Harmony Game as *n* increases. We identify three other large-*n* results, one of which shows the opposite: single-step advantages of cooperation may disappear as *n* grows. To facilitate comparisons, we define *A* throughout as the more cooperative type based on the single-step game, so that *a* > *d*. Thus, *AA* pairs survive better than *BB* pairs. We consider both infinite and finite populations but in neither case is there spatial or any other kind of structure. We do not develop a detailed biological, ecological or behavioral model. Akin to what is done in models of diploid viability selection, we describe the game and its results directly in terms of survival probabilities. Our results are applicable to a wide range of specific scenarios in which payoffs affect viability (as opposed to fertility or fecundity) and in which partnerships may either enhance survival or detract from it.

### 1.1. Evolutionary dynamics

Comparing *a*(*n*) to *c*(*n*) and *b*(*n*) to *d*(*n*) in Table 1 indicates whether the more cooperative strategy *A* yields the higher or the lower payoff given partners of type *A* and *B*, respectively. In game theory and much of evolutionary game theory which treats populations, these two comparisons are relevant because individuals choose or modify their strategies based on their knowledge of the game and its outcomes (Sandholm, 2010). When individuals cannot alter or choose their behaviors but payoffs affect fitness, the same sorts of evolutionary models are used to predict changes in the frequency of hardwired behaviors in a population. In this section, we briefly summarize the evolutionary game dynamics of infinite populations.

The replicator equation, in which general payoffs are interpreted as contributions to individual fitness (Taylor and Jonker, 1978; Hofbauer et al., 1979; Zeeman, 1980; Schuster and Sigmund, 1983; Hofbauer and Sigmund, 1988, 1998) provides an evolutionary setting for the classification of two-player games. In a survival game payoffs are fitnesses, specifically viabilities. If *x* is the relative frequency of *A* in an infinite population, the replicator equation gives the instantaneous rate of change

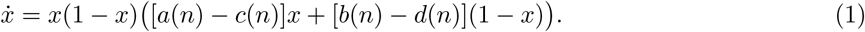

Equation (1) illustrates the evolutionary consequences of partner-dependent payoffs when pairs are formed at random in proportion to the frequencies of *A* and *B*. When *x* ≈ 1, so that *A* is very common and nearly all partners are *A*, it is the sign of *a*(*n*) − *c*(*n*) that determines whether *A* is favored in the population. On the other hand, when *x* ≈ 0, so that nearly all partners are *B*, the sign of *b*(*n*) − *d*(*n*) is what matters. Beyond this, Eq. (1) shows that for any value of *x*, it is the sign of [*a*(*n*) − *c*(*n*)]*x* + [*b*(*n*) − *d*(*n*)](1 − *x*) that determines whether *A* is favored. Thus, in a population and in evolution the relative magnitudes of both *a*(*n*) − *c*(*n*) and *b*(*n*) − *d*(*n*) are important, such that one may dominate the other at a given value of *x*. This is not something that can be gleaned directly from Table 1.

Without mutation, the fates of *A* and *B* in this infinite population are entirely determined by Eq. (1). From any starting point, *x* will move deterministically toward one of three possible equilibria which are the solutions of 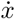 = 0 and which may correspond to the evolutionarily stable strategies mentioned in Section 1. Two monomorphic equilibria always exist, 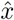 = 0 and 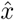 = 1. The polymorphic equilibrium

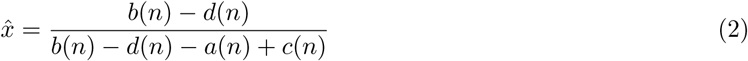

might also exist, depending on the payoffs. It exists, which is to say that Eq. (2) gives a biologically meaningful value, when *a*(*n*), *b*(*n*), *c*(*n*) and *d*(*n*) are such that 0 < 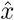 < 1.

Equation (1) defines the same four types of symmetric two-player games. In the PD case, *a*(*n*) − *c*(*n*) < 0 and *b*(*n*) − *d*(*n*) < 0, and Eq. (1) shows that 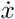 < 0 for all values of *x* ∈ (0,1). Starting at any value *x* < 1, the frequency of *A* will decrease to zero. The polymorphic equilibrium given by Eq. (2) does not exist. In the SH case, *a*(*n*) − *c*(*n*) > 0 and *b*(*n*) − *d*(*n*) < 0. Then *A* is disfavored when *x* is below the polymorphic equilibrium 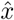 in Eq. (2) and favored when *x* > 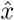. The polymorphic equilibrium exists and is unstable. In the HD case, *a*(*n*) − *c*(*n*) < 0 and *b*(*n*) − *d*(*n*) > 0. Then *A* is favored when *x* < 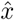, and disfavored when *x* > 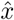. The polymorphic equilibrium exists and is stable. In the HG case, *a*(*n*) − *c*(*n*) > 0 and *b*(*n*) − *d*(*n*) > 0. Then 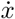 > 0 and *A* is favored for all *x* ∈ (0,1). The polymorphic equilibrium does not exist. Figure 1.1 shows examples of a Prisoner’s Dilemma, a Stag Hunt and a Hawk-Dove survival game, depicting payoffs in their contexts (upper panels) and associated shapes of 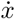 (lower panels). *A* Harmony Game is not shown but would be another case of directional selection, like Figure 1.1D but with 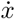 > 0 for all *x* ∈ (0,1).

**Figure 1:**
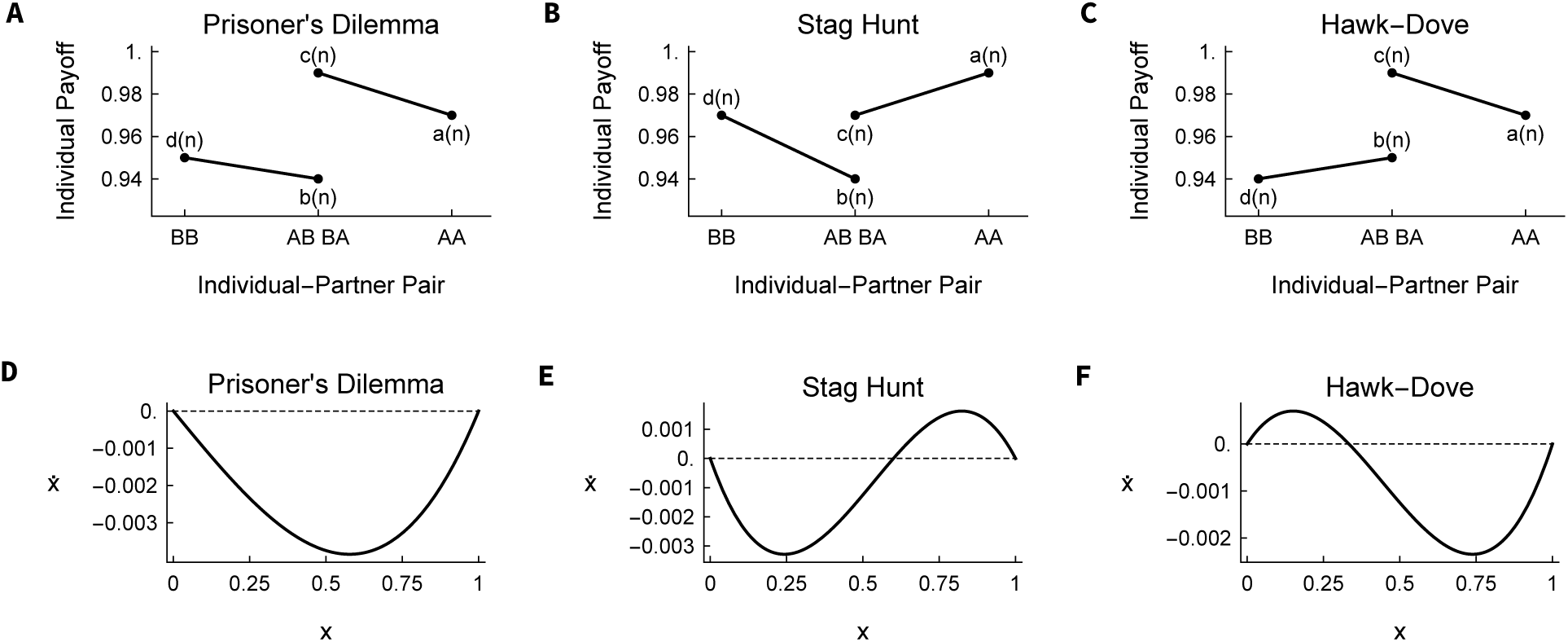
A and D: Prisoner’s Dilemma with payoffs *a* = 0.97, *b* = 0.94, *c* = 0.99, *d* = 0.95 (a linear transformation of the classic *R* =3, *S* = 0, *T* = 5, *P* = 1 of Axelrod (1984), e.g. *a* = 0.94 + *R*/100). B and E: Stag Hunt, with *a* = 0.99, *b* = 0.94, *c* = *d* = 0.97 (corresponding to a stag value of 0.05 and a hare value of 0.03 added to a baseline survival probability of 0.94). C and F: Hawk-Dove, with *a* = 0.97, *b* = 0.95, *c* = 0.99, *d* = 0.94 (corresponding to a cost of fighting of 0.03 and a resource value of 0.04, with a baseline survival probability of 0.95). Line segments in A, B and C connect the survival probabilities for *B* (left) versus *A* (right) when each occurs with given type of partner. The curves in D, E and F show 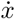 from Eq. (1). In diploid population genetics, these three cases for 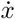 are called directional selection (D), underdominance (E) and overdominance (F).

These four types of games defined by the shape of 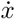 over the interval [0,1] are identical to what is observed in classical models of diploid viability selection (e.g. see section 4.2 in Nagylaki, 1992). The difference is that, barring atypical phenomena such as meiotic drive (Dunn, 1953; Sandler and Novitsky, 1957) or segregation-distortion (Sandler and Hiraizumi, 1960; Hartl, 1974), standard diploid models always have *b*(*n*) = *c*(*n*). It is by allowing *b*(*n*) ≠ *c*(*n*) that two-player games introduce the paradox of the Prisoner’s Dilemma, in which *c*(*n*) > *a*(*n*) and *d*(*n*) > *b*(*n*) but *a*(*n*) > *d*(*n*). In order to have *c*(*n*) > *a*(*n*) and *d*(*n*) > *b*(*n*) in a standard diploid model it would be necessary to have *a*(*n*) < *d*(*n*). Accordingly, the Prisoner’s Dilemma is the most stringent form of a cooperative dilemma (Doebeli and Hauert, 2005; Hauert et al., 2006; Nowak, 2012). A cooperative dilemma exists when (i) mutual cooperation results in a higher payoff than mutual non-cooperation, so that *a*(*n*) > *d*(*n*) when *A* represents cooperation, but (ii) there is incentive to be non-cooperative in at least one of three ways: (iia) *c*(*n*) > *a*(*n*), (iib) *d*(*n*) > *b*(*n*) or (iic) *c*(*n*) > *b*(*n*) (Nowak, 2012). The Prisoner’s Dilemma includes all three of these incentives. Games with fewer barriers to cooperation, such as the Stag Hunt and the Hawk-Dove game, represent relaxed cooperative dilemmas. The Harmony Game involves no cooperative dilemma and presents no barrier to cooperation.

## 2. An *n*-step survival game between two players

Our survival game always includes *n* iterations and begins with a pair of individuals. In each step, both might survive or one, the other or both might die. These same outcomes hold for the complete game, only the probabilities of surviving will be smaller. We consider two unconditional strategies, *A* and *B*. When both players are present, their individual survival probabilities are given by the single-step version of Table 1 with *a*(1) ≡ *a*, *b*(1) ≡ *b*, *c*(1) ≡ *c* and *d*(1) ≡ *d*. However, the next iteration must be faced by whoever has survived so far, until all *n* iterations are done. Thus, an individual might have to play alone. Then the survival probabilities become *a*_0_ for *A* without a partner and *d*_0_ for *B* without a partner. In all of what follows, *A* will be the more cooperative strategy, meaning that *a* > *d*.

The *n*-step survival game is a stochastic process with six possible states: the paired states *AA*, *AB* and *BB*, the loner states *A* and *B*, and the state in which both individuals have died which we denote Ø. It is convenient to represent a single iteration using the matrix

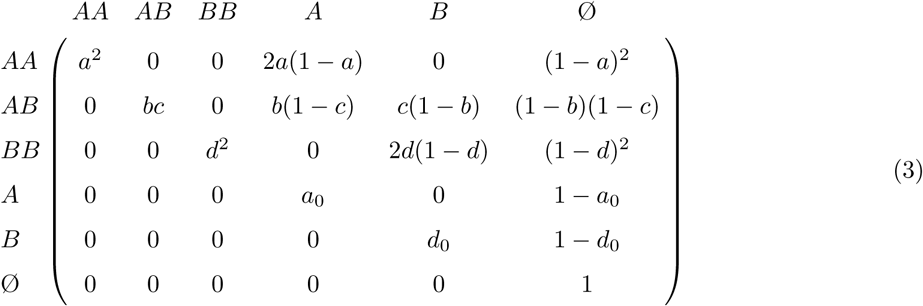

with entries equal to the transition probabilities among the six states. For example, the transition from state *AB* to state *B* means that the *B* individual survives and the *A* individual dies. In a single step, this occurs with probability *c*(1 − *b*). Note that the transitions in Eq. (3) include the fates of both individuals but the individuals are not labeled Individual and Partner as they are in Table 1.

The process described by Eq. (3) is depicted in Fig. 2. State Ø is an absorbing state. There is no possibility of transition between the paired states *AA*, *AB*, and *BB*. Instead each of these feeds either straight into state Ø, from which there is no escape, or into one of the loner states, *A* or *B*, and from there into Ø. Therefore, transitions from any of the starting, paired states to absorption in state Ø involve either one or two changes of state. Further, all of the transitions that cannot occur in a single iteration (the 0 entries in the matrix) cannot ever occur regardless of the number of iterations. Because of this simple structure, the *n*-step transition probabilities can be calculated directly by conditioning on the times these transitions take place, i.e. on their positions in the sequence of *n* iterations. It is also possible to compute the *n*-step transition probabilities using standard techniques for Markov chains, and this provides a useful framework for decomposing the process and describing its behavior when *n* is large.

**Figure 2:**
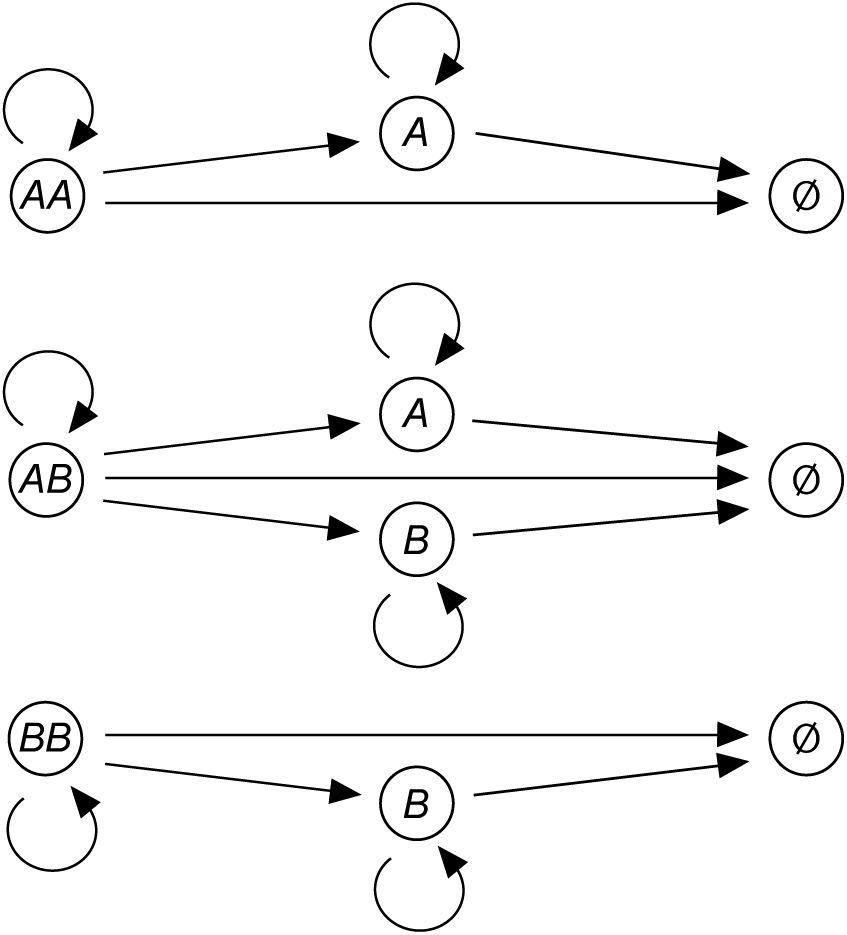
Flow diagram of the stochastic process given by the matrix of Eq. (3). Arrows show all possible transitions among the six states (*AA*, *AB*, *BB*, *A*, *B*, Ø). States *AA*, *AB*, *BB*, *A* and *B* are transient. State Ø is absorbing. The process always begins in one of the three states on the left, *AA*, *AB* or *BB*. After *n* iterations, it may be in any of the six possible states.

Details of the matrix approach are given in the Appendix, but two key features of it inform our presentation. The first is the fact of the absorbing state (Ø) in which both individuals have died. It will be reached eventually, meaning in the limit *n* → ∞. For large *n* the game process will be just the approach to this state. Second, when *n* is large, the rate of approach to state Ø will be given by the largest non-unit eigenvalue of the matrix in Eq. (3). Because it is an upper triangular matrix, the eigenvalues are simply the entries on the diagonal. By convention, we call the largest of these λ_1_ = 1 and note that this corresponds to the eventual
absorption in state Ø. In all, we have

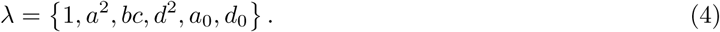

Following the discussion of Table 1 and Eq. 1, we expect the fates of *A* and *B* in this iterated game to depend on the relative magnitudes of *a* versus *c* and b versus *d*. Equation (4) suggests that their fates will also depend on the relative magnitudes of the pair survival probabilities, *a*^2^, *bc* and *d*^2^, and on the loner survival probabilities, *a*_0_ and *d*_0_, and that this dependence may be especially strong when *n* is large.

We compute the *n*-step pair survival probabilities directly as

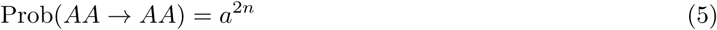

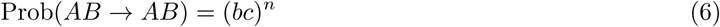

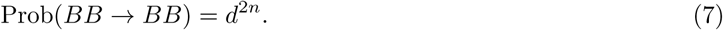

Then, by considering the possibility that one of the individuals might die in step 1 ≤ *i* ≤ *n* and the other individual survives to the end of the game, we have

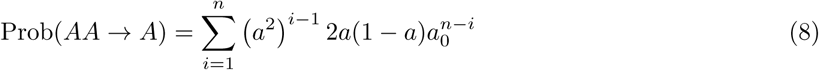

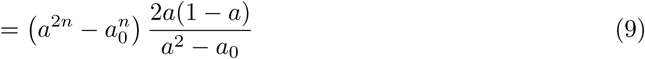

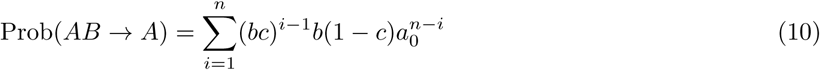

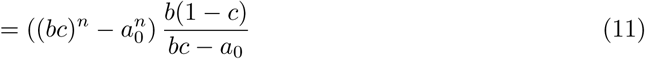

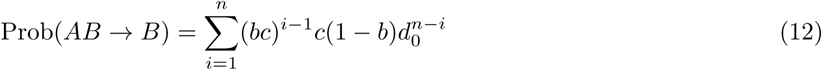

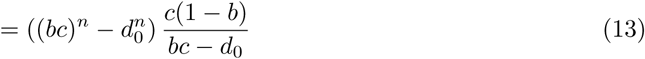

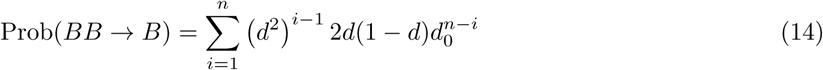

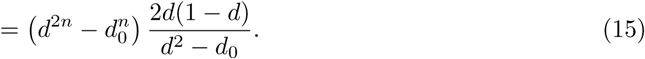

The only other possibility is that neither individual survives, so we also have

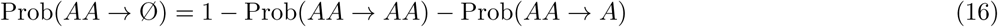

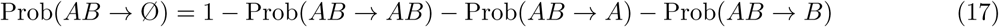

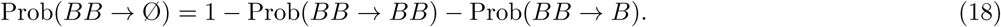

In the Appendix we show how these probabilities are obtained using techniques for Markov chains.

Equations (8) through (15) make it clear that this paired survival process is one in which the fortunes of individuals may change, perhaps drastically depending on the values of *a*_0_ and *d*_0_ relative to *a*, *b*, *c* and *d*. Consider starting state *AB*. Equation (12) shows how the probability of being in state *B* at the end of *n* iterations is computed. In words, both individuals survive for some time (*i* − 1 steps), then *A* dies, and *B* survives the rest of time (*n* − *i* steps). If *A* dies, *B* trades its individual survival probability of *c*, which *B* enjoys in the presence and *A*, for a loner survival probability of *d*_0_. It could be that *d*_0_ < *c*, making *B* worse off after *A* dies. In fact, *B*’s fate is closely tied with *A*’s because this switch could occur quickly if *A*’s survival probability, *b*, is small. Of course, while there is a cost to *B* when *A* dies (assuming *d*_0_ < *c*), the death of *A* in this partnership also represents a rather direct disadvantage to *A*. In order to understand whether *A* or *B* will prevail in evolution, it is necessary to account for the full dynamics of reproduction in a population, with fitnesses are determined by this game. We will take this up in Section 3.

In the *n*-step game, the differences *a*(*n*) − *c*(*n*) and *b*(*n*) − *d*(*n*) give the conditions under which *A* is favored. The *n*-step payoffs, or survival probabilities, are computed by accounting for the two ways an individual may survive the game. An individual survives if both it and its partner survive or if it survives but its partner dies. The total probabilities of individual survival in each kind of partnership are

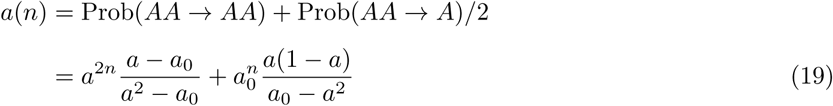

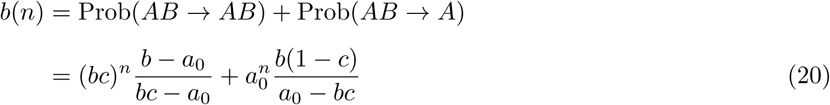

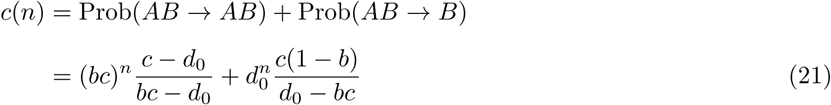

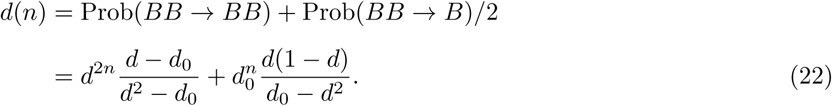

The transitions *AA* → *A* and *BB* → *B* are adjusted by a factor of 1/2 because they are equally likely to happen by the death of the partner as by the death of the focal individual. Equations (19) through (22) are our general results, namely the *n*-step survival probabilities for individuals of type *A* and *B* given each kind of initial partnership. They are exact for any values of *a*, *b*, *c*, *d*, *a*_0_ and *d*_0_ greater than zero and less than one, and for any number of iterations *n* ≥ 1. As expected, when *n* =1, they reduce to the single-step survival probabilities *a*(1) = *a*, *b*(1) = *b*, *c*(1) = *c* and *d*(1) = *d*.

The two key differences in payoff are then

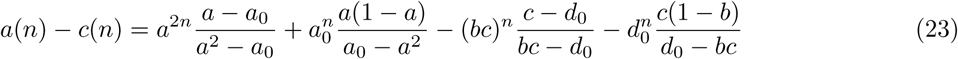

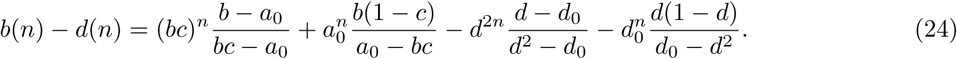

Each term in Eqs. (23) and (24) includes a factor 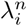 for one of the non-unit eigenvalues (*a*^2^, *bc*, *d*^2^, *a*_0_, *d*_0_). These factors are positive. The denominator of each term is the difference between two eigenvalues. If we know the eigenvalues, we know whether the denominators are positive or negative. In some cases the numerator might be negative, but by the definition of *a*, *b*, *c*, *d*, *a*_0_ and *d*_0_ as probabilities, the numerator will be positive if the denominator is positive. Thus Eqs. (23) and (24) are written so that the sign of each term indicates whether it favors *A* (+) or disfavors *A* (−) when the denominator is positive. We do this to facilitate the analysis of large *n*, in which case *a*(*n*) − *c*(*n*) and *b*(*n*) − *d*(*n*) will come to be dominated by the terms involving the largest non-unit eigenvalue.

Another way to write Eqs. (23) and (24) is

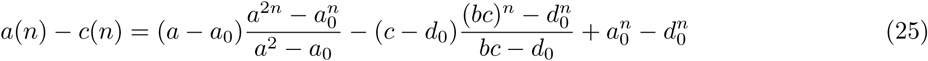

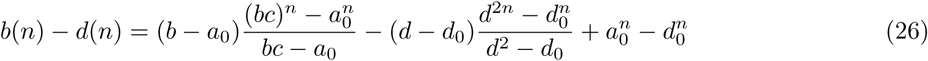

which brings attention to the possible survival value of having a partner and of being *A* versus *B* without a partner. Thus, *a* − *a*_0_ and *c* − *d*_0_ in Eq. (25) are the increments to the survival probabilities of *A* and *B* when each has partner *A* compared to when they are alone. Likewise, *b* − *a*_0_ and *d* − *d*_0_ in Eq. (26) are the corresponding increments for *A* and *B* with partner *B* versus alone. Note that the fractions multiplying these increments are always positive. All else being equal, *a*(*n*) − *c*(*n*) increases with *a* − *a*_0_ > 0 but decreases with *c* − *d*_0_ > 0, and *b*(*n*) − *d*(*n*) increases with *b* − *a*_0_ > 0 but decreases with *d* − *d*_0_ > 0. Negative increments have the opposite effects. For example, if *B* suffers by having a *B* partner compared to being alone, then *d* − *d*_0_ < 0, which in turn favors *A* via the difference *b*(*n*) − *d*(*n*). The last two terms of both Eqs. (25) and (26) quantify the *n*-step effects of differences in the loner survival probabilities, *a*_0_ and *d*_0_. If *a*_0_ > *d*_0_ the contribution to both *a*(*n*) − *c*(*n*) and *b*(*n*) − *d*(*n*) is positive, and if *a*_0_ < *d*_0_ it is negative.

These five possible increments to individual survival, namely *a* − *a*_0_, *b* − *a*_0_, *c* − *d*_0_, *d* − *d*_0_ and *a*_0_ − *d*_0_, in Eqs. (25) and (26) are helpful for understanding some of our results and the structure of survival games generally. However, it is also clear from the additional presence of the pair-state eigenvalues in Eqs. (25) and (26) as well as in Eqs. (23) and (24), that these five increments do not entirely determine the prospects for cooperation. The relative magnitudes of the five eigenvalues will also be important. This may be seen in the denominators in Eqs. (25), (26), (23) and (24), which contrast the survival of pairs to the survival of loners: *a*^2^ − *a*_0_, *bc* − *a*_0_, *bc* − *d*_0_, *d*^2^ − *d*_0_. The preceding expressions, Eqs. (23) and (24), are especially useful for predicting the structure of the game when *n* is large.

## 3. Four types of prolonged survival games

Here, we present analytical results for large *n* and use our exact results to illustrate how the prospects for *A* depend on *n* for intermediate values. The analysis of large *n* allows us to answer questions such as: Is there a number of iterations beyond which *A* is unambiguously favored? More generally, we ask how the game might change for *A*, for better or for worse, as the number of iterations increases. We present results both for *a*(*n*) − *c*(*n*) and *b*(*n*) − *d*(*n*) and in terms of the unified predictions of Eq. (1).

Equation (1) is a standard replicator equation for a symmetric two-player game. In the Appendix, we show how it may be derived for the *n*-step survival game. This has two notable features: the game works by removing individuals from the population instead of affecting their rates of reproduction and the derivation follows pairs of individuals rather than single individuals. Also in the Appendix, we take a discrete-time, population-genetic approach to show that the change in frequency of *A* over one generation is

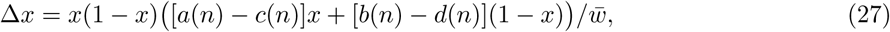

which is identical in form to 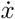 in Eq. (1) but scaled by the average fitness, *w*. Equation (27) is directly comparable to classical results for diploid viability selection. Importantly, *w* > 0. Therefore, all of the conclusions concerning the ultimate fate of the cooperative type *A* in the population, based on the sign of the change in frequency, will be the same whether one appeals to Eq. (1) or Eq. (27).

Each of the *n*-step payoffs *a*(*n*), *b*(*n*), *c*(*n*) and *d*(*n*) depends on two of the non-unit eigenvalues, and each eigenvalue is raised to the power *n*. Thus *a*(*n*), *b*(*n*), *c*(*n*) and *d*(*n*) all decrease to zero as *n* tends to infinity. Surviving more steps is less likely than surviving fewer steps. As a consequence, the two key fitness differences, *a*(*n*) − *c*(*n*) and *b*(*n*) − *d*(*n*), and the instantaneous rate of change, 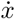, also approach zero in the limit *n* → ∞. We study the approach to this neutral limit in order to understand whether increasing *n* fundamentally alters the prospects for cooperation. For example, if 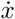 < 0 when *n* =1 but the neutral limit 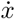 = 0 is approached from above, then there exists a value of *n* beyond which 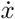 > 0. In this case, the single-step game favors the non-cooperative type *B* but large-*n* games favor the cooperative type *A*.

In the following subsections, we describe four main categories of large-*n*, or prolonged, survival games. These are characterized both in the usual way by the relative values of *a*, *b*, *c* and *d*, that is by the structure of the single-step game, and by the rank order of non-unit eigenvalues *a*^2^, *bc*, *d*^2^, *a*_0_ and *d*_0_. We have defined the cooperative strategy A throughout by *a* > *d*, which guarantees that *a*^2^ > *d*^2^. Thus *d*^2^ will never be the largest non-unit eigenvalue. The first two types of prolonged survival games we describe are those for which *a*^2^ is the largest and those for which *bc* is the largest. The third and fourth types of prolonged survival games are ones in which lone individuals survive with high probability. Simplifications emerge when *n* is large because the *n*-step payoffs become dominated by their largest terms. We present approximations for large *n*, showing just the leading terms of *a*(*n*) − *c*(*n*) and *b*(*n*) − *d*(*n*), and of 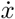.

### 3.1. Games in which cooperation will prevail

The first case we consider is when having a partner significantly enhances survival and it is the *AA* type not the *AB* type that survives best. This is case in which *a*^2^ is the largest non-unit eigenvalue: *a*^2^ > *bc*, *a*_0_, *d*_0_. Thus, the probability that both members of an *AA* pair survive a single iteration is greater than the corresponding probability for any other pair and for either type of lone individual.

The condition *a*^2^ > *bc* may be true of any kind of two-player game, but the chance it is depends on the game. One way to quantify this is to consider the total parameter space of two-player, single-step survival games. Because *a*, *b*, *c* and *d* are probabilities and here it is assumed that *a* > *d*, this space is equal to half of a four-dimensional hypercube. Further, since *a* > *d* implies *a*^2^ > *d*^2^, there are just three possible relations of the pair-state eigenvalues, *a*^2^, *bc* and *d*^2^. Either *bc* > *a*^2^, *a*^2^ > *bc* > *d*^2^ or *d*^2^ > *bc*. The latter two satisfy the condition *a*^2^ > *bc*. Table 2 lists the proportions of the total parameter space satisfying the criteria for each type of single-step game. In addition, Table 2 shows the percentages of the game-specific parameter regions corresponding to each of the three possible relations of the pair-state eigenvalues. So, for example, 90% of PD-type games have *a*^2^ > *bc*, compared to 20% of HD-type games.

**Table 2:**
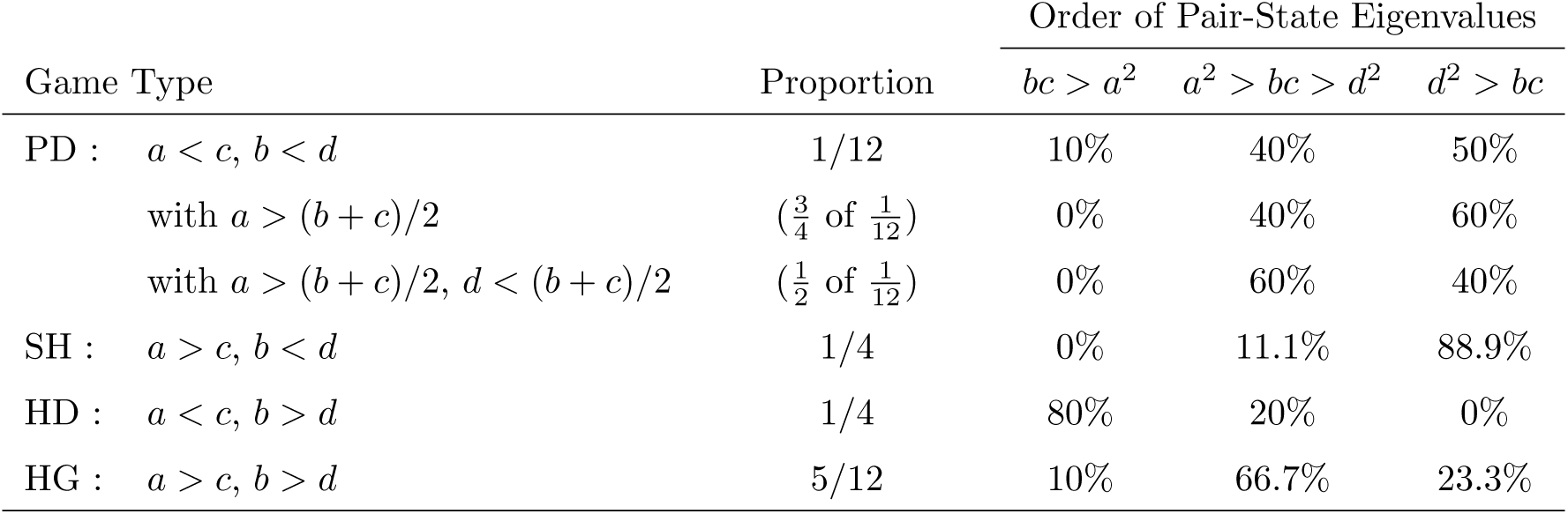
Partitioning of the four-dimensional hypercube into fractions meeting the game conditions in the first column. The types of single-step games are abbreviated PD, SH, HD and HG for Prisoner’s Dilemma, Stag Hunt, Hawk-Dove game, and Harmony Game. Exact results were obtained by integration in Mathematica (Wolfram Research, Inc.). Percentages are rounded to the nearest 0.1%. Note that in all cases *a* > *d* by definition, so it is always true that *a*^2^ > *d*^2^.

| Game Type | Proportion | Order of Pair-State Eigenvalues |  |  |
| --- | --- | --- | --- | --- |
| | | $bc > a^2$ | $a^2 > bc > d^2$ | $d^2 > bc$ |
| PD : $a < c, b < d$ | 1/12 | 10% | 40% | 50% |
| with $a > (b + c)/2$ | ($\frac{3}{4}$ of $\frac{1}{12}$) | 0% | 40% | 60% |
| with $a > (b + c)/2, d < (b + c)/2$ | ($\frac{1}{2}$ of $\frac{1}{12}$) | 0% | 60% | 40% |
| SH : $a > c, b < d$ | 1/4 | 0% | 11.1% | 88.9% |
| HD : $a < c, b > d$ | 1/4 | 80% | 20% | 0% |
| HG : $a > c, b > d$ | 5/12 | 10% | 66.7% | 23.3% |

When the single-step game is a canonical Prisoner’s Dilemma, it is always true that *a*^2^ > *bc*. This is due to the assumption that the reward payoff must be larger than the average of the temptation payoff and the sucker’s payoff (see, e.g. Rapoport and Chammah, 1965; Axelrod, 1984), in which case we have

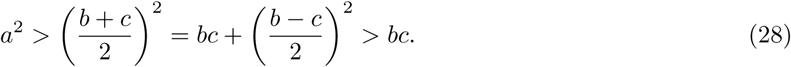

In what follows, it will be important to know which is the second-largest non-unit eigenvalue, especially in the consideration of finite populations in Section 4. Thus, the two cases *a*^2^ > *bc* > *d*^2^ and *d*^2^ > *bc* are distinguished in Table 2. We find that canonical Prisoner’s Dilemmas are 40% *a*^2^ > *bc* > *d*^2^ and 60% *d*^2^ > *bc*. If we also require that the punishment payoff must be less than the average of the temptation payoff and the sucker’s payoff (e.g. see p. 219 of Cressman, 2005), this 40: 60 split is reversed.

It would be possible to expand Table 2 by including *a*_0_ and *d*_0_ and integrating over the six-dimensional space of all parameters. There would be more than three relations among the five non-unit eigenvalues. We do not pursue this here. For the prolonged games considered in this section and in Section 3.2, it is taken for granted that neither *a*_0_ nor *d*_0_ is the largest non-unit eigenvalue. In contrast, in Sections 3.3 and 3.4 it is taken for granted that one of *a*_0_ or *d*_0_ is the largest non-unit eigenvalue or they both are and they are equal. If we were to include *a*_0_ and *d*_0_ in Table 2, we expect they would often be larger than *a*^2^ and *bc* due to the fact that these pair-state eigenvalues are products of probabilities.

There is just one term in Eqs. (19) through (22) which includes the factor *a*^2*n*^, corresponding to what is assumed here to be the largest non-unit eigenvalue, *a*^2^. It is in the expression for *a*(*n*). Thus, we have

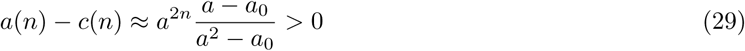

when *n* is very large. Equation (29) gives the approximate value of *a*(*n*) − *c*(*n*) as it approaches zero in the limit *n* → ∞. This result shows that whatever value or sign *a*(1) − *c*(1) = *a* − *c* might have in a single iteration, there exists a value of *n* above which *a*(*n*) − *c*(*n*) will be greater than zero.

The leading term in the other key difference, *b*(*n*) − *d*(*n*), will depend on which of *bc*, *d*^2^, *a*_0_ or *d*_0_ is the next-largest eigenvalue. Consider, for example, a game which strongly enhances survival, such that *a*^2^, *bc*, *d*^2^ > *a*_0_, *d*_0_. If *bc* is the next largest eigenvalue after *a*^2^, then from Eq. (24) we have

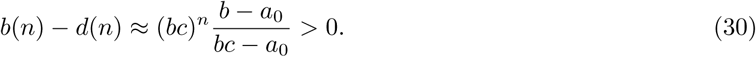

In this case, the large-*n* game becomes a Harmony Game in which *A* is unambiguously favored. On the other hand, if the second largest eigenvalue is *d*^2^, then from Eq. (24) we have

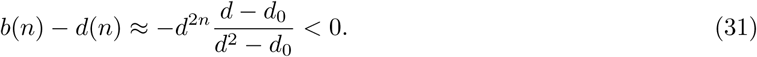

In this case, the large-*n* game becomes or remains a Stag Hunt game, so *A* does not become unambiguously favored. The chances of being in either of these two sub-cases are given in Table 2. Similar arguments can be applied when *a*_0_ or *d*_0_ is the second-largest non-unit eigenvalue. Then Eq. (24) shows that for large *n*, *b*(*n*) − *d*(*n*) > 0 if *a*_0_ is the second-largest, and *b*(*n*) − *d*(*n*) < 0 if *d*_0_ is the second-largest.

Regardless of which is the second-largest non-unit eigenvalue, if *a*^2^ is the largest then *a*(*n*) − *c*(*n*) will be much greater than *b*(*n*) − *d*(*n*) when *n* is large. This may be expressed as

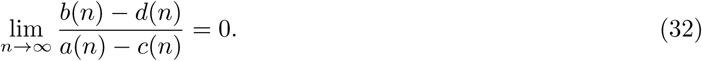

Thus, when *n* is very large, the advantage in survival that *A* has over *B* when the partner is *A* will be much greater than whatever difference there is in survival between *A* and *B* when the partner is *B*. One may argue from an evolutionary standpoint that the large-*n* Stag Hunt game, with *a*(*n*) − *c*(*n*) in Eq. (29) and *b*(*n*) − *d*(*n*) in Eq. (31), will be a very relaxed cooperative dilemma. Specifically, owing to Eq. (32), a large-*n* Stag Hunt of this sort will have its unstable polymorphic equilibrium 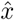 in Eq. (2) close to zero, with the result that *A* will be favored over most of the range of *x*.

Finally, when *n* is very large the replicator equation, Eq. (1), becomes

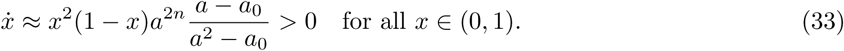

We conclude that, if *n* is large enough in the case of this first type of prolonged survival game, the more cooperative type *A* will be uniformly favored in an infinite population. This result requires only that *a*^2^ is the largest non-unit eigenvalue. It is robust to differences in the loner survival probabilities *a*_0_ and *d*_0_. As long as *a*^2^ is large enough, the non-cooperative type *B* cannot be rescued by an advantage in loner survivability. Table 2 shows that *a*^2^ > *bc* for the great majority of possible single-step survival games, the exception being games of the Hawk-Dove type, of which only 20% will have this property.

Figure 3 illustrates how the two key differences in payoff, *a*(*n*) − *c*(*n*) and *b*(*n*) − *d*(*n*), change as *n* increases for two examples based on the Prisoner’s Dilemma. In Fig. 3A, the survival probabilities for individuals with partners are *a* = 0.8, *b* = 0.6, *c* = 0.9, *d* = 0.7. Thus, the cooperative type *A* incurs an 11% loss of fitness compared to *B* when the partner is *A* and a 14% loss when the partner is *B*. The survival probabilities for loners are much smaller and identical: *a*_0_ = *d*_0_ = 0.3. The eigenvalues are *a*^2^ = 0.64, *bc* = 0.54, *d*^2^ = 0.49, *a*_0_ = 0.3 and *d*_0_ = 0.3. This game starts as a Prisoner’s Dilemma (PD) when *n* =1. For intermediate numbers of iterations, specifically *n* = 3 and *n* = 4, the game changes to one in which *A* is disfavored when rare (*b*(*n*) − *d*(*n*) < 0) but favored when common (*a*(*n*) − *c*(*n*) > 0), as in the Stag Hunt (SH). Then as *n* grows it becomes a game in which *A* is uniformly favored, as in the Harmony Game (HG).

**Figure 3:**
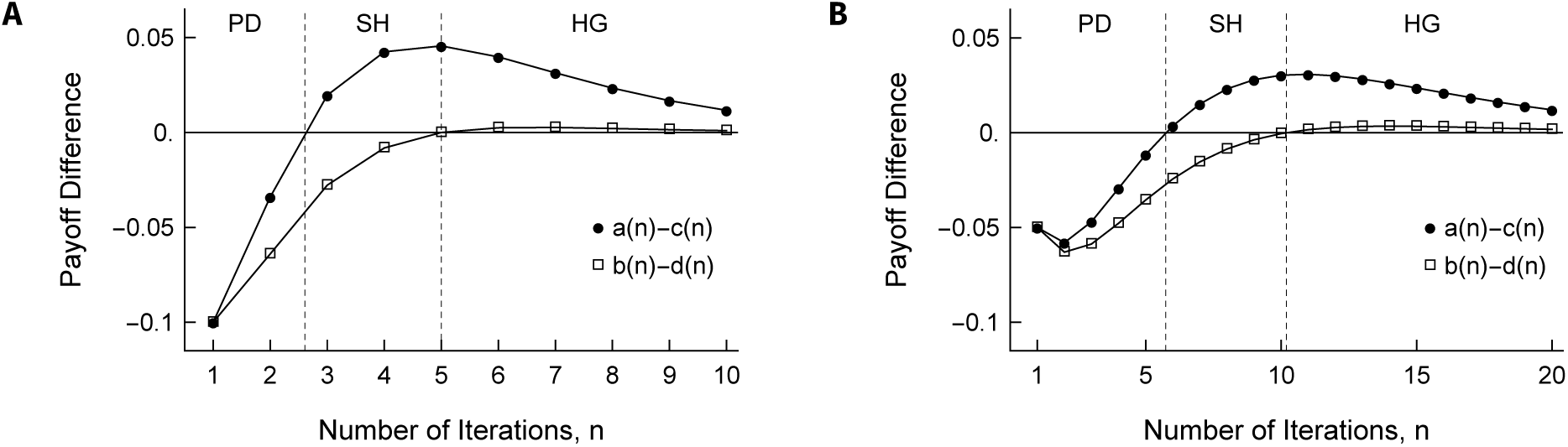
Values of the two key differences, *a*(*n*) − *c*(*n*) and *b*(*n*) − *d*(*n*), as a function of the number of iterations, for two different survival games which are Prisoner’s Dilemmas when *n* =1. Panel A: *a* = 0.8, *b* = 0.6, *c* = 0.9, *d* = 0.7 and *a*_0_ = *d*_0_ = 0.3. Panel B: *a* = 0.9, *b* = 0.8, *c* = 0.95, *d* = 0.85 and *a*_0_ = *d*_0_ = 0.65.

Figure 3B displays results for a very similar game, but one in which the single-iteration probabilities of individual survival are shifted closer to 1 by a factor of 1/2. That is, *a* = 0.9, *b* = 0.8, *c* = 0.95, *d* = 0.85 and *a*_0_ = *d*_0_ = 0.65. Now the eigenvalues are *a*^2^ = 0.81, *bc* = 0.76, *d*^2^ = 0.7225, *a*_0_ = 0.65 and *d*_0_ = 0.65. Overall, the picture is similar to Figure 3A, with the three phases PD then SH then HG. But now in moving from *n* = 1 to *n* = 2, the selection against *A* becomes stronger rather than weaker. This is more in line with the usual notion of iterated games, in which payoffs are assumed to accrue additively. In survival games, however, this decrease cannot continue because all games become neutral as *n* approaches infinity.

Figure 3 may be compared to Fig. 4 (Model 2.A.III) of De Jaegher and Hoyer (2016), with their number of attacks being analogous to *n*. A point of contrast between our model based on multiplying single-step survival probabilities and the behavioral and ecological scenario, Model 2 of De Jaegher and Hoyer (2016) is that in our model the payoff differences *a*(*n*) − *c*(*n*) and *b*(*n*) − *d*(*n*) may be non-monotonic, as they are in Fig. 3, whereas these are always monotonic in Model 2 of De Jaegher and Hoyer (2016). We will return to Model 2 of De Jaegher and Hoyer (2016) and make further comparisons in the Discussion.

**Figure 4:**
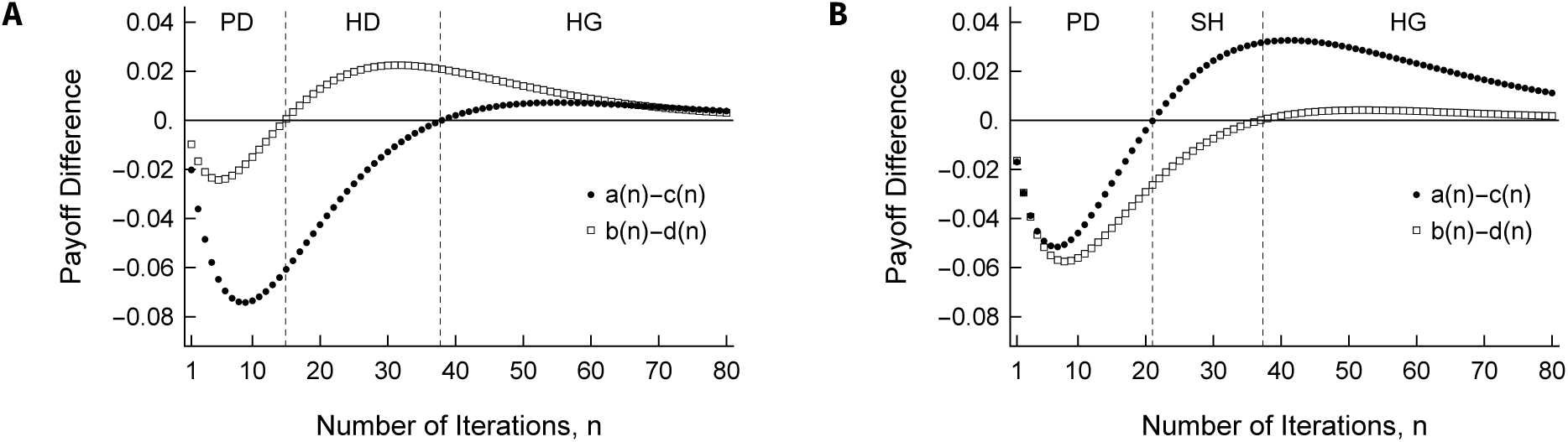
Values of *a*(*n*) − *c*(*n*) and *b*(*n*) − *d*(*n*) over *n* = 1… 80 iterations for two games which begin as Prisoner’s Dilemmas at *n* = 1. In both cases, the loner survival probabilities are *a*_0_ = *d*_0_ = 0.9. Panel A: *a* = 0.97, *b* = 0.94, *c* = 0.99, *d* = 0.95; these are the same as in Fig. 1.1A and D. Panel B: *a* = 0.9733, *b* = 0.94, *c* = 0.99, *d* = 0.9567; this differs from panel *A* only in that *a* and *d* are spaced evenly between *c* and *b*.

Because survival probabilities decrease with each iteration, *n* is a measure of the adversity faced by individuals, or the harshness of the environment. Another way to measure adversity is directly in terms of single-step survival probabilities. Adversity is greater when *a*, *b*, *c*, *d*, *a*_0_ and *d*_0_ are smaller. Figure 3 provides an example of how an increase in this sort of environmental challenge affects the outcome of the game. Single-step survival probabilities are smaller in Fig. 3A, so we would say that the environment is harsher in Fig. 3A than in Fig. 3B. Comparing the two shows that increasing this kind of adversity causes the cooperative type *A* to become unambiguously favored (HG) sooner, that is for smaller *n*.

Figure 4 shows how *a*(*n*) − *c*(*n*) and *b*(*n*) − *d*(*n*) change as the number of iterations increases for two Prisoner’s Dilemmas with survival probabilities even closer to 1. In Fig. 4A, the game at *n* = 1 is identical to the example in Fig. 1.1A and 1.1D. Again, the payoffs in this game follow the classical spacing (*R* = 3, *S* = 0, *T* = 5, *P* = 1) of Axelrod (1984). For comparison, the game in Fig. 4B has the same value of the temptation payoff (*c* = 0.99) and the sucker payoff (*b* = 0.94) but has reward *a* and punishment *d* spaced evenly between these, as was the case in Fig. 3. In both Fig. 4A and 4B, the loner survival probabilities are *a*_0_ = *d*_0_ = 0.9. Both show the phenomenon of a worsening dilemma (greater disadvantage for *A*) at small *n*, then a switch to a relaxed dilemma and finally to a Harmony Game. In Fig. 4B, the intermediate, relaxed dilemma is a Stag Hunt game, as in Fig. 3. In Fig. 4A, the intermediate, relaxed dilemma is instead a Hawk-Dove game. Equations (25) and (26) are helpful in understanding the difference between Fig. 4A and Fig. 4B. In moving from Fig. 4A to Fig. 4B, the increment *a* − *a*_0_ in Eq. (25) is boosted by 0.0033 and the increment *d* − *d*_0_ in Eq. (26) is boosted by 0.0067. This increases *a*(*n*) − *c*(*n*) and decreases *b*(*n*) − *d*(*n*).

Figure 5 shows how *a*(*n*) − *c*(*n*) and *b*(*n*) − *d*(*n*) change as *n* increases for examples of the Stag Hunt and the Hawk-Dove game. Parameters are as in Fig. 1.1 with the addition of *a*_0_ = *d*_0_ = 0.9 for both models. As with the examples in Fig. 4, these are cases in which *a*^2^ is the largest non-unit eigenvalue, so Eqs. (29), (30)

**Figure 5:**
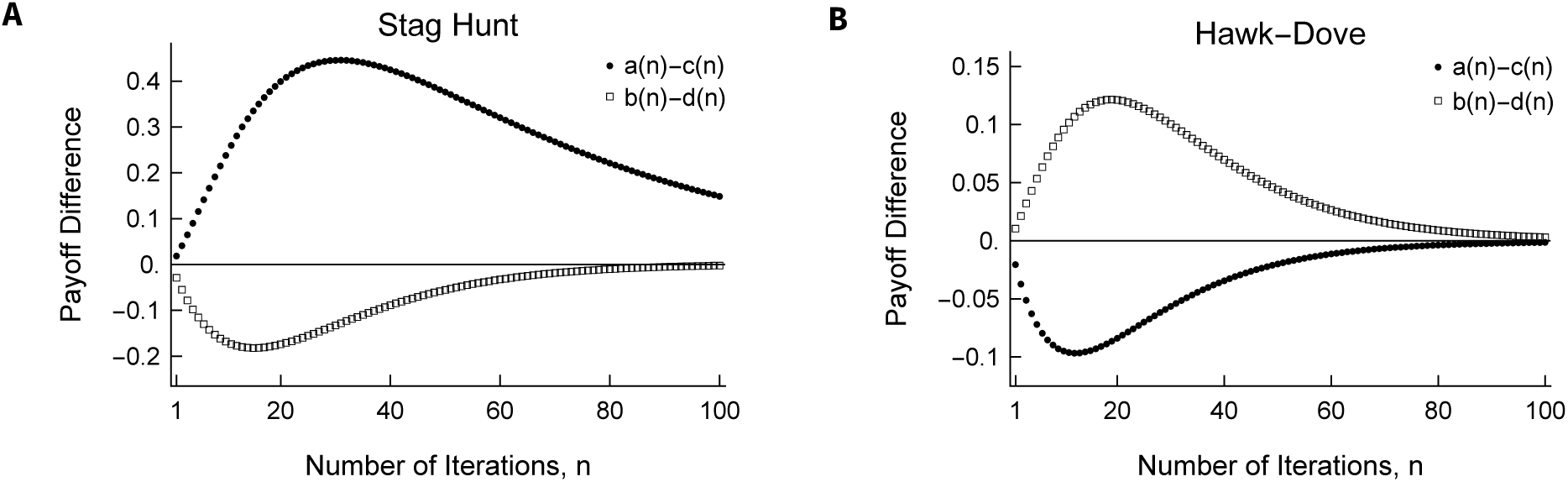
Values of *a*(*n*) − *c*(*n*) and *b*(*n*) − *d*(*n*) over *n* = 1… 100 iterations for two games which begin as a Stag Hunt and as a Hawk-Dove game at *n* = 1. In both cases, as in Fig. 4, the loner survival probabilities are *a*_0_ = *d*_0_ = 0.9. Panel A: a = 0.99, *b* = 0.94, *c* = *d* = 0.97; these are the same as in Fig. 1.1B and E. Panel B: a = 0.97, *b* = 0.95, *c* = 0.99, *d* = 0.94; these are the same as in Fig. 1.1C and F.

and (31) apply. In contrast to what is seen in Fig. 4, one or the other of *a*(*n*) − *c*(*n*) and *b*(*n*) − *d*(*n*) is still negative in Fig. 5 even at *n* = 100. From *n* = 1 to beyond *n* = 100, these remain a Stag Hunt and a Hawk-Dove game. In the Stag Hunt of Fig. 5A, the next-largest eigenvalue is *d*^2^, so *b*(*n*) − *d*(*n*) will remain negative. However, as can already be seen in the figure, it will be much smaller in magnitude than *a*(*n*) − *c*(*n*). In the Hawk-Dove game of Fig. 5B, the next-largest eigenvalue is *bc*, so *b*(*n*) − *d*(*n*) will remain positive. But since *a*^2^ is the largest non-unit eigenvalue, *a*(*n*) − *c*(*n*) will become positive (when *n* = 615, from numerical solution) and eventually will become much larger than *b*(*n*) − *d*(*n*).

Another way to view the results presented in Figs. 3, 4 and 5 is in terms of 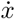 rather than *a*(*n*) − *c*(*n*) and *b*(*n*) − *d*(*n*). Figure 6 shows this for the examples of Fig. 4. Now the overall shape of 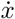 can be seen as it moves from *n* =1, where it is a Prisoner’s Dilemma identical to what is depicted in Fig. 1.1D, to *n* = 60, where it is a Harmony Game. In both panels, the thick contour line traces 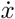 = 0, showing the polymorphic equilibrium in Eq. (2) when it exists. In Fig. 6A, this equilibrium enters at *x* = 0 and exits at *x* = 1. In between, the shape of 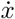 is like that of Fig. 1.1F, namely similar to the Hawk-Dove game in which *A* is favored when rare and disfavored when common. In Fig. 6B, the equilibrium enters at *x* =1 and exits at *x* = 0. Here the shape of 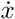 for intermediate *n* is like that of Fig. 1.1E, namely similar to the Stag Hunt in which *A* is disfavored when rare and favored when common. Note that although *a*(*n*) − *c*(*n*) and *b*(*n*) − *d*(*n*) vary non-monotonically in these cases (Fig. 4), the equilibrium 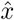 is monotonic in *n*.

**Figure 6:**
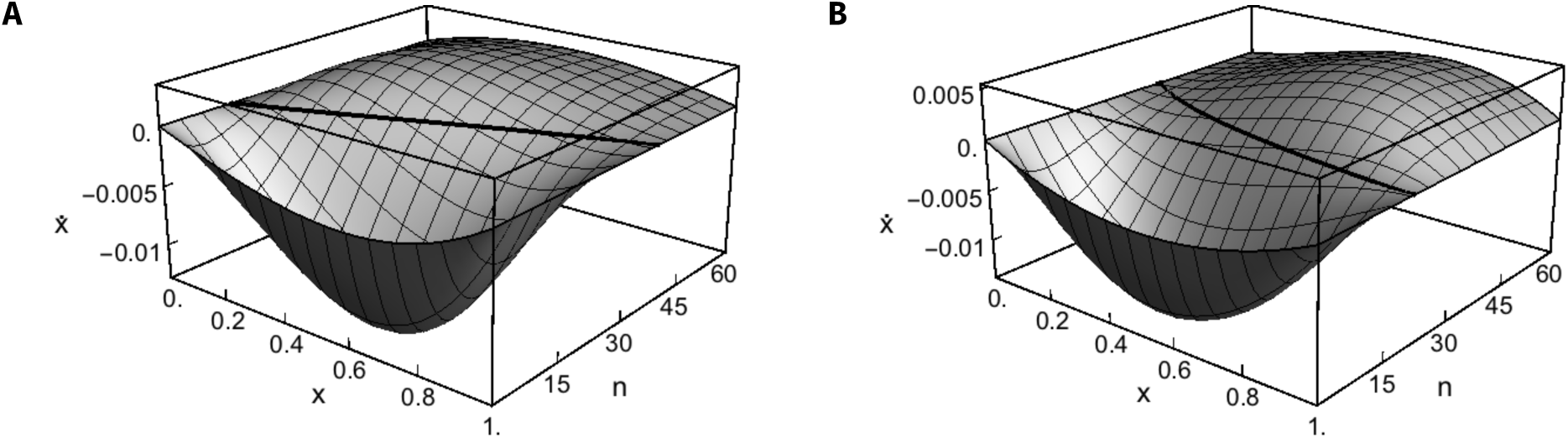
Plots of 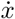 for the models in Fig. 4, that is for two different Prisoner’s Dilemmas at *n* = 1. Panel A: *a* = 0.97, *b* = 0.94, *c* = 0.99, *d* = 0.95 and *a*_0_ = *d*_0_ = 0.9. Panel B: *a* = 0.9733, *b* = 0.94, *c* = 0.99, *d* = 0.9567 and *a*_0_ = *d*_0_ = 0.9. In both panels, the thick contour line is drawn at 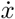 = 0, which corresponds to the polymorphic equilibrium 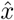 of Eq. (2) for each value of *n*. For simplicity of presentation, we plot the surface as a smooth function of *n*, whereas in reality it is discrete.

Figure 7 shows how the shape of 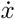 changes as the number of iterations increases for the Stag Hunt and the Hawk-Dove game depicted in Fig. 5. Note that to aid in visualizing these surfaces, the viewpoints are different in the two panels and the ranges of *n* are different than in Fig. 5. As with the Prisoner’s Dilemmas in Figs. 3 and 6, these are both cases in which *a*^2^ is the largest non-unit eigenvalue. Following the discussion of Fig. 5, although the polymorphic equilibria traced by the thick contour lines are still in the picture for the largest values of *n* in Fig. 7, in both cases *A* will become favored if *n* is large enough. In the Stag Hunt of Fig. 7A or 5A, 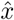 → 0 as *n* → ∞. In the Hawk-Dove game of Fig. 7B or 5B, 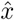 will exit the biologically relevant interval and become larger than one when *a*(*n*) − *c*(*n*) first becomes positive, when *n* = 615.

**Figure 7:**
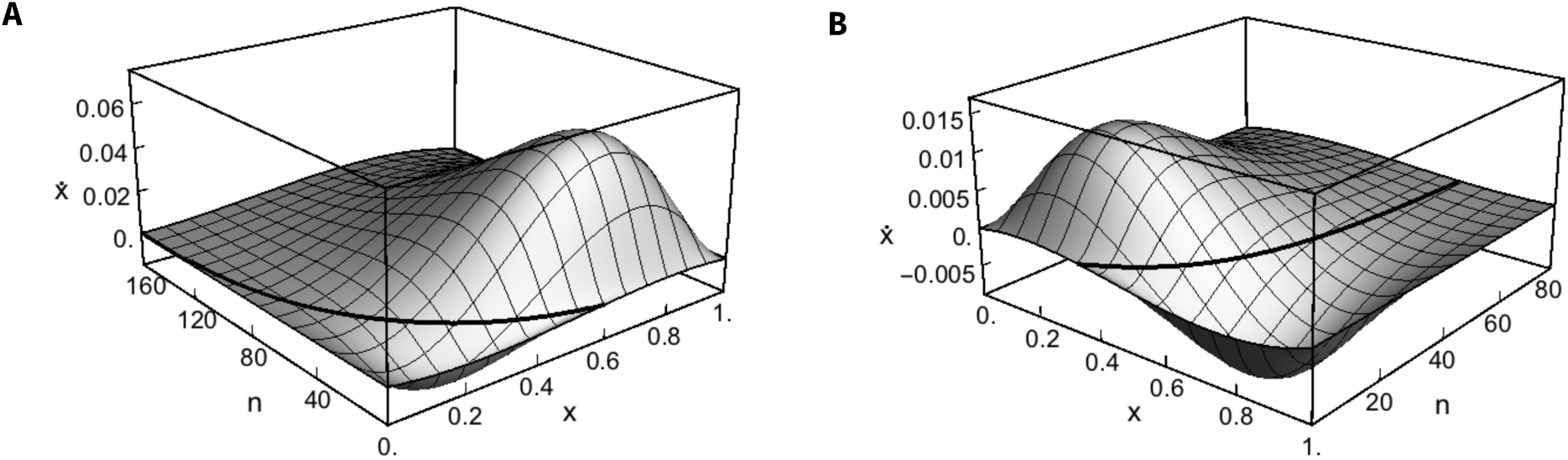
Plots of 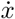 for the models in Fig. 5. For the *n* = 1, Stag Hunt in panel A: *a* = 0.99, *b* = 0.94, *c* = *d* = 0.97 and *a*_0_ = *d*_0_ = 0.9. For the *n* = 1, Hawk-Dove game in panel B: *a* = 0.97, *b* = 0.95, *c* = 0.99, *d* = 0.94 and *a*_0_ = *d*_0_ = 0.9. In both panels, the thick contour line is drawn at 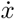 = 0, which corresponds to the polymorphic equilibrium 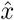 of Eq. (2) for each value of *n*. Note that the viewpoints and the ranges of *n* for the two panels are different.

### 3.2. Games which preserve non-cooperative behaviors

Here we consider the case that the game enhances survival significantly and it is *AB* that fares best. Thus, *bc* is the largest non-unit eigenvalue. Table 2 shows that this will occur most readily when the single-step game is a Hawk-Dove game. It might also occur for Prisoner’s-Dilemma type games, but only if the canonical assumption *a* > (*b* + *c*)/2 is violated. It will not occur when the single-step game is a Stag Hunt. Here, again, we assume that having a partner significantly enhances survival, so *bc* > *a*_0_, *d*_0_.

There are two terms in Eqs. (19) through (22) which include the factor (*bc*)^*n*^. They are in the expressions for *b*(*n*) and *c*(*n*). For very large *n* we have

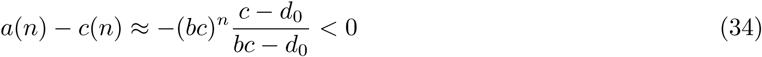

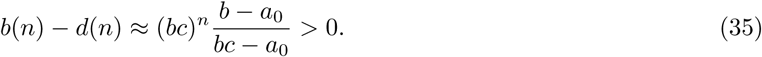

Thus, survival games of this sort either stay or become Hawk-Dove games as *n* increases. In addition, *a*(*n*) − *c*(*n*) and *b*(*n*) − *d*(*n*) will be of the same order of magnitude as *n* tends to infinity. In the previous section, where *AA* survived best, increasing *n* converted single-step Prisoner’s Dilemmas, Stag Hunts and some Hawk-Dove games into progressively more relaxed cooperative dilemmas until, eventually, *A* became favored. Here, Eqs. (34) and (35) show that single-iteration Hawk-Dove games in which *AB* survives best are robust to such alteration. Equations (34) and (35) also show that if a survival game is a non-canonical Prisoner’s Dilemma at *n* =1, such that *bc* rather than *a*^2^ is largest non-unit eigenvalue, the *n*-step game will eventually become a Hawk-Dove game rather than a Harmony Game.

When *bc* is the largest non-unit eigenvalue, the large-*n* approximation for the change in frequency is

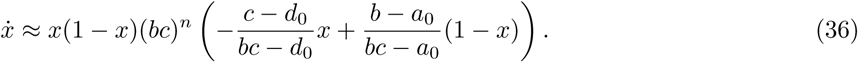

We deduce that, for large *n*, 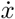 > 0 if *x* < 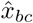 and 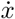 < 0 if *x* > 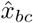, where

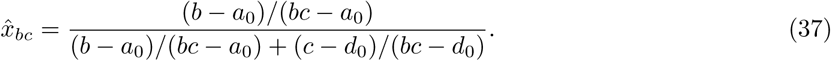

This frequency cutoff may be interpreted as the probability, when *n* is large, that if *AB* plays the game and only one individual survives, it is the *A* individual. For reference, if *a*_0_ and *d*_0_ are both very small, then Eq. (37) reduces to b/(b + *c*). If the population frequency of *A* is smaller than this probability in Eq. (37), then *A* is favored, otherwise it is disfavored. This cutoff is of course equal to the limiting (*n* → ∞) value of Eq. (2) when bc is the largest non-unit eigenvalue, which is stable in this case.

Figure 8 depicts both how *a*(*n*) − *c*(*n*) and *b*(*n*) − *d*(*n*) change and how 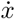 changes as *n* increases for a Hawk-Dove example in this case. It may be more easily seen in Fig. 8B that this example remains a Hawk-Dove game with a stable polymorphic equilibrium (the thick contour line) as *n* increases, even as the game first intensifies for intermediate *n* then approaches neutrality for large *n*.

**Figure 8:**
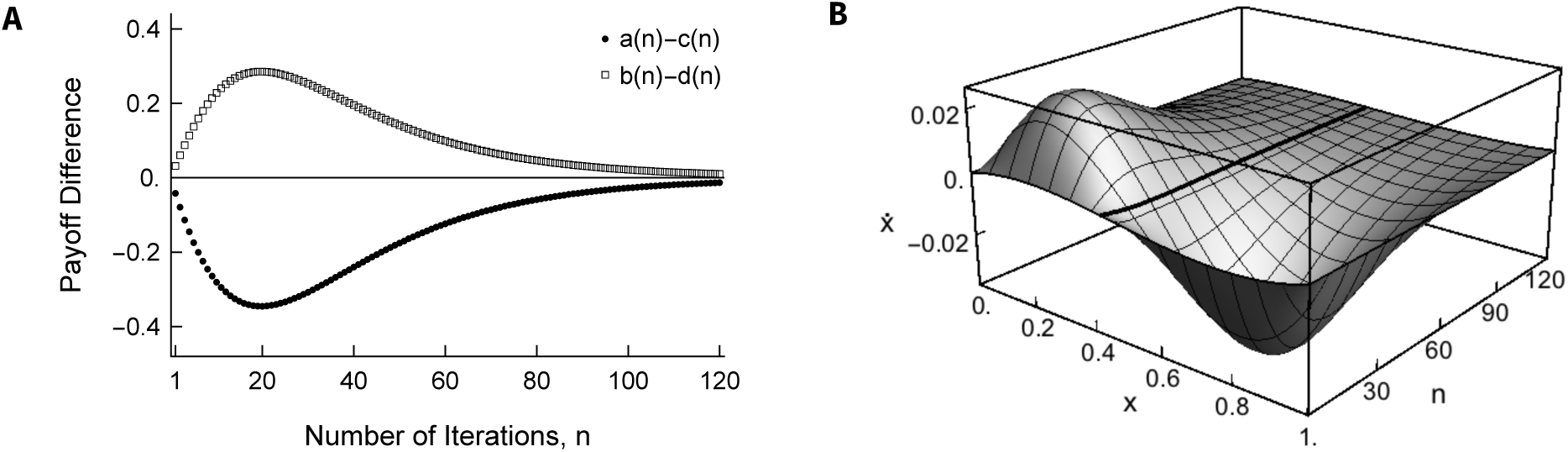
Hawk-Dove type game in which *bc* is the largest eigenvalue: payoffs *a* = 0.95, *b* = 0.97, *c* = 0.99, *d* = 0.94 and *a*_0_ = *d*_0_ = 0.9. The thick contour line in *B* shows 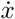 = 0, which stabilizes to 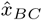 = 0.5625 as *n* increases.

### 3.3. Games ruled by loner survivability

In both of the previous sections, it was assumed that having a partner provided a substantial boost to survival. In this section and the next, we consider cases in which having a partner does not enhance survival. If either a lone *A* or a lone *B* has the highest probability of surviving each iteration of the game, then either *a*_0_ or *d*_0_ is the largest non-unit eigenvalue. Two terms in Eqs. (19) through (22) include the factor 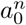—they are in the expressions for *a*(*n*) and *b*(*n*)—and two terms include the factor 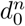—they are in the expressions for *c*(*n*) and *d*(*n*). If *a*_0_ is the largest non-unit eigenvalue, then for very large *n* we have

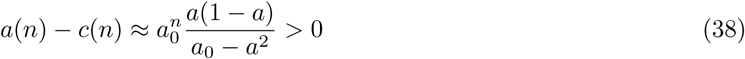

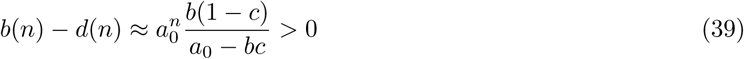

and if *d*_0_ is the largest non-unit eigenvalue, we have

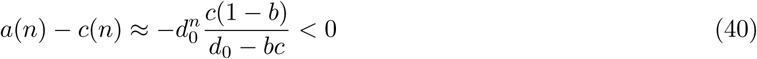

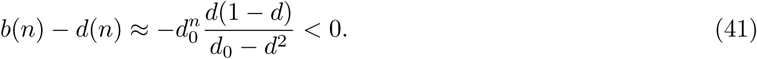

Thus, if single individuals of one type have a higher probability of survival than single individuals of the other type and than any pair of individuals, that type will become favored in the population as the number of iterations grows, regardless of the structure of pairwise interactions in the single-step game. The corresponding approximations for *x* can be inferred from Eqs. (38) through (41).

### 3.4. Games with a pairwise survival deficit

A related but probably more interesting case is when single individuals have higher survival probabilities than any pair of individuals but the loner survival probabilities are identical. Then having a partner does not enhance survival but the two-player game may still provide an advantage to one strategy or the other. In this case *a*_0_ = *d*_0_ is the largest non-unit eigenvalue. When *n* is very large, using Eqs. (23) and (24) and substituting *a*_0_ for *d*_0_, we have

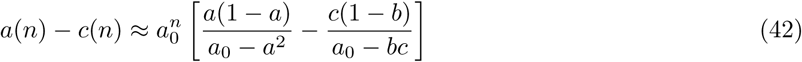

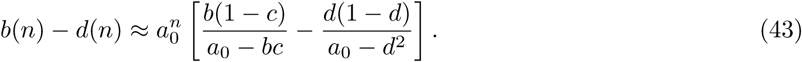

Whether these differences are greater or less than zero will depend on which of the two terms in the brackets is larger in absolute value. This situation, in which the game is played starting in the pair state with a survival deficit, is similar in spirit to the way the Prisoner’s Dilemma was originally formulated, with two individuals in custody, each wanting to get out. Here, as in Section 3.3, we do not show the corresponding result for 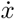, which can be inferred from Eqs. (42) and (43).

Consider the case of a strong deficit, specifically with *a*_0_ = do not too small and all of *a*, *b*, *c* and *d* close to zero. Then, neglecting terms of *a*^2^, *bc* and *d*^2^ which will be even smaller, Eqs. (42) and (43) become

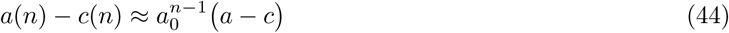

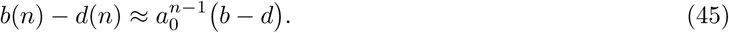

Thus, with a strong deficit, the results for large *n* will be of the same type as the results for *n* = 1.

When the deficit is milder, no simple approximations for Eqs. (42) and (43) are available, but numerical analysis shows that the conclusions of Section 3.1 can be reversed. Figure 9 shows two examples of how *a*(*n*) − *c*(*n*) and *b*(*n*) − *d*(*n*) may change as *n* increases, converting a single-step Harmony Game and a single-step Hawk-Dove game into Prisoner’s Dilemmas. In both examples, *a*_0_ = *d*_0_ is the largest non-unit eigenvalue. In Fig. 9A, the game becomes a Stag Hunt for intermediate values of *n*. In Fig. 9B, the game changes directly from a Hawk-Dove game into a Prisoner’s Dilemma as *n* increases. Thus, when having a partner does not significantly enhance survival but does cause payoff differences—and is the only source of these differences—there is no expectation that the prospects for cooperation will improve as the number of iterations increases. These cooperative dilemmas may intensify rather than becoming more relaxed.

**Figure 9:**
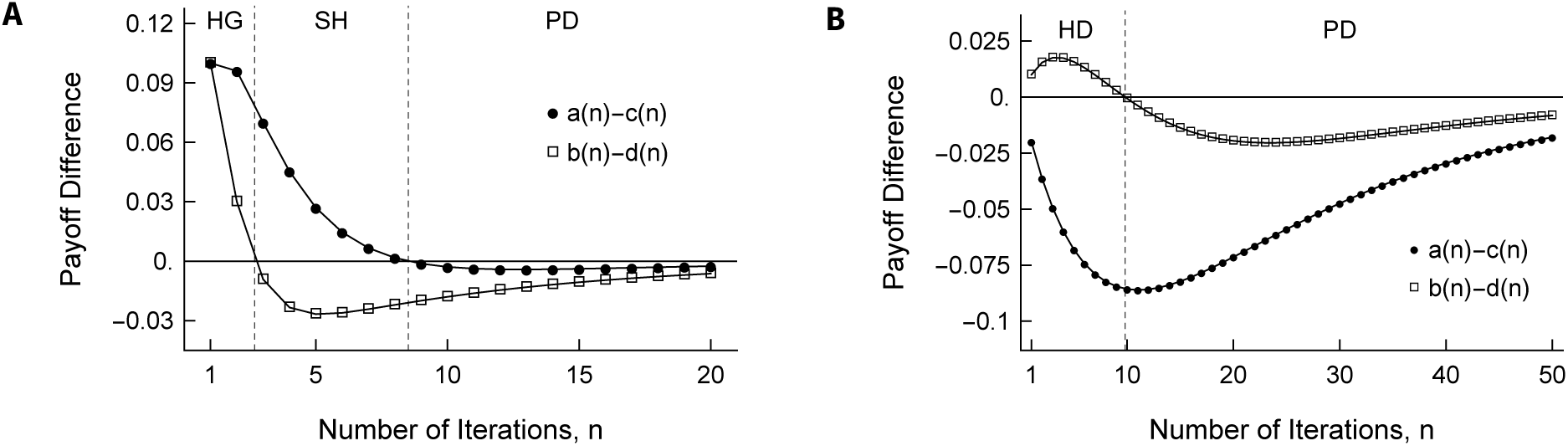
Values of *a*(*n*) − *c*(*n*) and *b*(*n*) − *d*(*n*) as *n* increases for two games which begin as a Harmony Game and as a Hawk-Dove game at *n* =1. Panel A: *a* = 0.8, *b* = 0.5, *c* = 0.7, *d* = 0.4 and *a*_0_ = *d*_0_ = 0.9. Panel B: Hawk-Dove *a* = 0.93, *b* = 0.91, *c* = 0.95, *d* = 0.90 and *a*_0_ = *d*_0_ = 0.95.

## 4. Evolutionary dynamics in a finite population

All populations are finite, so changes in the frequency of *A* will be stochastic rather than deterministic. Here we describe these changes for a particular population model and check the validity of our previous conclusions based on the deterministic predictions for 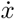 (or Δ*x*). In a finite population, *x* is discrete, not continuous. If *N* is the population size and *K* is the number of *A* individuals, then *K* ∈ (0,1,…, *N* − 1, *N*) and *x* ∈ (0,1/*N*,…, (*N* − 1)/*N*, 1). We study the process of jumps in *K* or *x* when both death and reproduction are stochastic. We continue to assume that the survival game is the only source of fitness differences and that the population is well-mixed in the sense that partners are chosen at random.

Due to the relative simplicity of its transitions and the ease with which the survival game may be embedded within it, we base our model on the haploid population-genetic model of Moran (1958). The standard version of the Moran model pairs single births and deaths such that the population size remains constant. In the absence of selection, each individual has probability 1/*N* of being chosen to reproduce in a given time step, and a 1/*N* chance to die. The individual chosen to reproduce makes an offspring which replaces the individual chosen to die. When selection is included, it usually acts by modifying the probability an individual is chosen, now based on its fitness, either to reproduce or to die.

We, instead, consider a model in which two randomly chosen individuals play the survival game. Players are chosen without replacement from the population. This precludes self-interaction, which is appropriate because we wish to model two individuals being sequestered from the rest of the population for a period of time. When the *n* iterations of the game are finished, both individuals may be alive, one may be alive or both may be dead. This is the only source of death in the population and the only place in the life cycle where selection acts. In each time step, these 0, 1 or 2 deaths are compensated by the same number of births. All individuals have an equal chance to reproduce, including the two who play the game. Thus, parents are sampled at random with replacement from the population. Finally, offspring inherit their parent’s type without modification, i.e. there is no mutation.

Let *K* be the current number of *A* individuals. After one time step, the number is *K*′, which will differ from *K* by at most 2 and is bounded by 0 and *n*. The numbers of *A* individuals in a series of time steps forms a Markov chain with an (*N* +1) × (*N* +1), pentadiagonal transition probability matrix. Because there is no mutation, the states *K* = 0 and *K* = *N* are absorbing. The transition probabilities are computed as products of three terms: one for the sampling of a pair of individuals to play the game, one for the outcome of the game and one for the sampling of parents. In all, we have

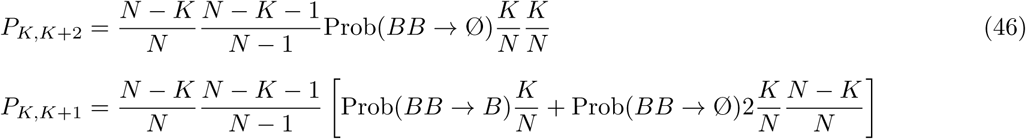

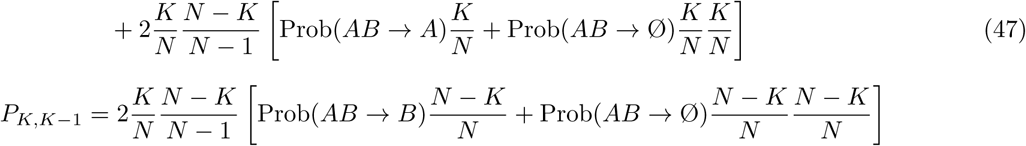

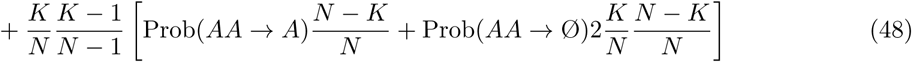

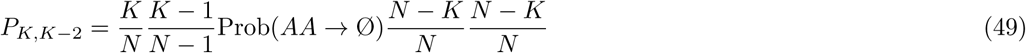

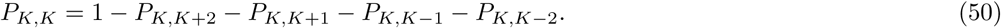

Although this process is not as simple as the standard Moran process, for which a number of exact results are available (Moran, 1962), as a pentadiagonal matrix it may still be amenable to study. Following our previous analysis of 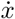, we focus on the change in the number of cooperators, Δ*K* = *K*′ − *K*, in particular on the sign and magnitude of *E*[Δ*K*] which measures the direction and strength of selection.

The expected value of Δ*K* is given by

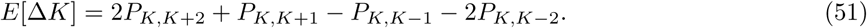

Using Eqs. (46) though (50) and simplifying yields

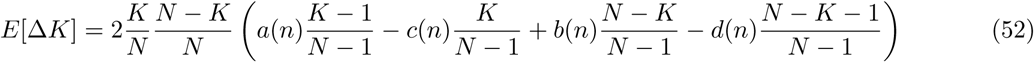

with *a*(*n*), *b*(*n*), *c*(*n*), *d*(*n*) as before in Eqs. (19) through (22). Because *x* = *K*/*N*, we also have *E*[Δ*x*] = *E*[Δ*K*]/*N*. Aside from the factor of 2, which arises because two individual are chosen to play the game and possibly die, Eq. (52) shows nearly the same dependence on *a*(*n*), *b*(*n*), *c*(*n*) and *d*(*n*) as the deterministic result for 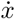 in Eq. (1). We define the selection coefficient

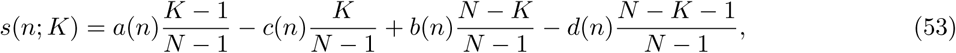

which captures the dependence of *E*[Δ*K*] on *a*(*n*), *b*(*n*), *c*(*n*) and *d*(*n*) in its entirety. Of course, the sign of *s*(*n*; *K*) determines the sign of *E*[Δ*K*] and *E*[Δ*x*].

Comparing *s*(*n*; *K*) in Eq. (53) to its deterministic counterpart in Eq. (1) reveals that when *N* is small or when *K* is small regardless of *N* the deterministic prediction is incorrect. The error comes from overestimating the importance of *AA* and *BB* pairings in the infinite-population model by implicitly allowing self-interaction. Our aim here is to reassess the conclusions of the previous section in light of such differences and especially in light of the fact that *A* can be fixed or lost from a finite population regardless of the value of *E* [Δ*K*]. Criteria for *A* being favored in a finite population are generally based on the probability it reaches fixation starting at *K* =1 versus the probability it is lost starting at *K* = *N* − 1 (Nowak et al., 2004; Lessard, 2005). If the probability of fixation of *A* is greater than the neutral probability 1/*N* and the probability of loss is less than 1/*N*, we might conclude that *A* is favored. When selection is weak, a number of analytical results for fixation probabilities and times in evolutionary games are available (Wu et al., 2010). Alternatively, in models with mutation the decision as to whether *A* is favored can be made by comparing the expected equilibrium frequency of *A* under neutrality versus selection (Rousset and Billiard, 2000). When the mutation rate is small, these two approaches become identical.

In Section 3, we used 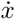 > 0 for all *x* ∈ (0,1) as a criterion for *A* being unambiguously favored. The analogue here is *E*[Δ*x*] > 0 for all *x* ∈ (1/*N*,…, (*N* − 1)/*N*). We take this to be a more stringent criterion than the one based on fixation probabilities. However, from Eqs. (52) and (53), it is clear that even when *AA* survives best and *a*^2^ is the largest non-unit eigenvalue, it will not necessarily be true that *E*[Δ*x*] > 0 for all *x* ∈ (1/*N*,…, (*N* − 1)/*N*). In particular, for the two terminal frequencies, we have

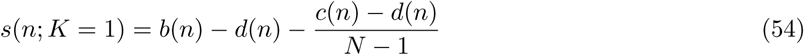

and

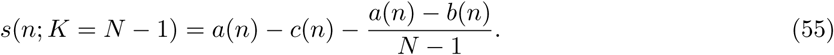

When there is only a single *A* individual, the possible benefit of the *AA* pair is unrealized. At the other extreme, when there is only one *B* individual, it is the *BB* pair that does not matter.

Therefore, even when *a*^2^ is the largest non-unit eigenvalue, it will always be possible to find a population size small enough that *A* remains disfavored even as *n* tends to infinity. In particular, if the population consists of just two individuals, there is only one polymorphic frequency, *K* = *N* - *K* = 1, and we have

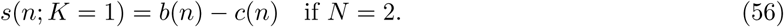

From Eqs. (20) and (21) we have

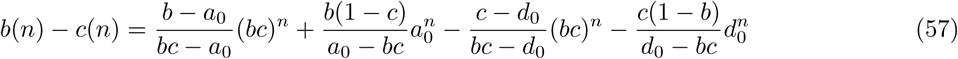

which will not in general be favorable to *A*. To illustrate, if we make the simplifying assumption that the loner fitnesses are the same (*a*_0_ = *d*_0_), then Eq. (57) reduces to

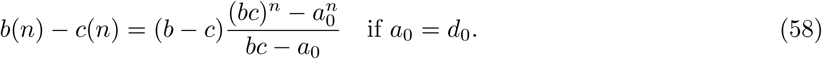

When *n* =1, this is simply *b* − *c*. Then as *n* becomes large, regardless of which eigenvalue (*bc* or *a*_0_ = *d*_0_) is larger, the sign of *b*(*n*) − *c*(*n*) will be the same as the sign of *b* − *c*. One of the ways in which a game may be considered a cooperative dilemma is that *b* < *c* (Nowak, 2012). Note that *b* < *c* in all of our numerical examples. When *N* = 2, none of the conclusions about relaxed cooperative dilemmas based on *x* and increasing *n* will hold. Just the opposite: *A* will be disfavored.

While there is no expectation that conclusions from an infinite-population model will be accurate for small populations, we expect their heuristic value to hold for moderately large *n*. In what follows, we focus on *N* ≫ 1. We begin as before by noting that our finite-population model collapses to a neutral Moran model in the limit *n* → ∞, although it would be one in which pairs of individuals rather than single individuals are chosen to die and to reproduce. Again, we answer the question about *A* being favored by studying the approach to this neutral limit. In our finite-population model, this neutral limit is *E*[Δ*x*] = 0. Then if, for example, *E*[Δ*x*] < 0 when *n* =1 but the neutral limit *E*[Δ*x*] = 0 is approached from above, then there exists a value of *n* above which *E*[Δ*x*] > 0. We briefly reconsider each type of prolonged game.

### Case I, reconsidered

This is the case in which *a*^2^ is the largest non-unit eigenvalue and cooperation is expected to prevail. The finite-population result corresponding to Eq. (33) is

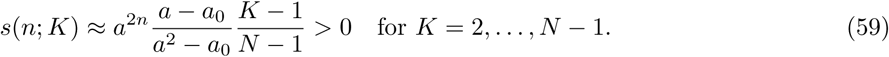

Thus, for *n* large enough, *E*[Δ*x*] > 0 for every frequency *x* = *K*/*N* except for one. We must investigate *K* =1 because, for a given large value of *n*, even though *a*(*n*) may dominate all other terms when *K* = 2,…, *N* − 1, it is only *b*(*n*), *c*(*n*) and *d*(*n*) that appear in *s*(*n*; *K* = 1) in Eq. (54). When *n* is large the sign and magnitude of *s*(*n*; *K* = 1) will be determined by the second largest non-unit eigenvalue. We consider the same two possibilities as before, namely when either *bc* or *d*^2^ is next-largest eigenvalue, and we continue to assume that the loner survival probabilities, *a*_0_ and *d*_0_ are small by comparison.

If the next largest eigenvalue is *bc*, then for *K* = 1 we have

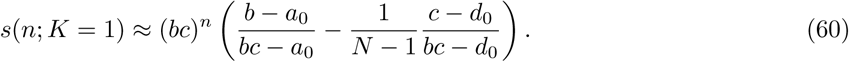

For very small *N* it could be that Eq. (60) is negative, but it will be positive for any moderate to large *N*, making *E*[Δ*x*] > 0 for *x* = 1/*N*. Combining this with Eq. (59), we conclude that here, with *n*^2^ > *bc* >
*d*^2^, *a*_0_, *d*_0_, if *n* is large enough, then *E*[Δ*x*] > 0 for all *K* = 1,…, *N* − 1. Previous Figs. 3 and 4 are examples of this case in which *A* becomes unambiguously favored.

If the second-largest non-unit eigenvalue is *d*^2^, we have

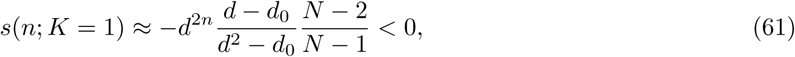

and *E*[Δ*x*] < 0 when *x* = 1/*N*. But, again, for large *n* it will also be true that *d*^2*n*^ ~ *a*^2*n*^, so the magnitude of the selection coefficient against *A* when *K* =1 will be small compared to selection coefficients in favor of *A* for every other value of K. We state without proof that in this case, if *n* is large enough, the probability of fixation of *A* starting from a single copy should be greater than the neutral probability 1/*N*, and the probability of loss of *A* starting from *N* − 1 copies should be less than 1/*N*. We conclude that *A* should be considered favored in this case as well.

Situations in which one of the loner survival probabilities, *a*_0_ or *d*_0_, is the second largest eigenvalue can be treated similarly. We do not pursue this in detail, but in Case III below we consider the possibility that one of these is the largest non-unit eigenvalue, and from that we can infer that nothing beyond the two possibilities above for *s*(*n*; *K* = 1) will arise. That is, as *n* grows, *s*(*n*; *K* = 1) will be either positive or negative but will be very small compared to *s*(*n*; *K*) which is positive for *K* = 2,…, *N* − 1. Thus, the conclusion that *A* becomes favored when *a*^2^ is the largest non-unit eigenvalue still holds.

### Case II, reconsidered

This is the case in which *bc* is the largest non-unit eigenvalue and the game tends to preserve the less cooperative strategy. The finite-population result corresponding to Eq. (36) is

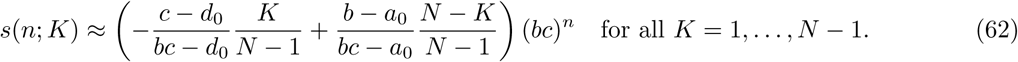

Thus, the terminal frequencies *x* = 1/*N* and *x* = 1 − 1/*N* do not require a separate treatment as they did under Case I. Here, just as in the infinite-population model, increasing *n* will not fundamentally alter the game. Selection pressure will tend to keep both *A* and *B* in the population.

### Case III, reconsidered

This is the case in which *a*_0_ or *d*_0_ is the largest non-unit eigenvalue. For these games ruled by loner survivability, we have

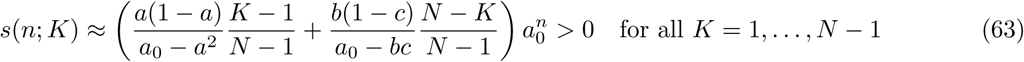

if *a*_0_ is the largest non-unit eigenvalue, and

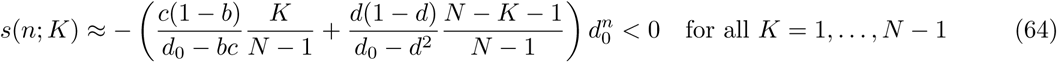

if *d*_0_ is the largest non-unit eigenvalue. The conclusions are unchanged in the modified Moran model relative to the infinite-population model. Whichever type survives best alone becomes favored when *n* is large.

### Case IV, reconsidered

This is the case in which *a*_0_ = *d*_0_ is the largest non-unit eigenvalue. For these games with a pairwise survival deficit, we have

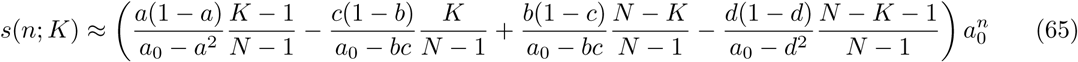

in general, and

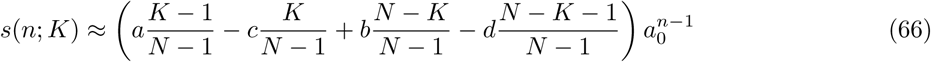

when the deficit is strong. If *K* =1, the first terms in the parentheses in Eqs. (65) and (66) disappear, making *A* worse off when it is very rare, because those terms are positive. When *K* = *N* − 1, the last terms in the parentheses in Eqs. (65) and (66) disappear, making *A* better off when *B* is very rare, because those terms are negative. It is not expected that this will improve the already dim the prospects for the evolution of cooperative behaviors in this case in which there is no survival advantage to having a partner.

## 5. Discussion

We have proposed a simple but general, biologically motivated survival game. It is a repeated game which always starts with a pair of individuals and includes a fixed number of iterations. Probabilities of survival are its expected payoffs. We have assumed that multi-step survival is a Markov process, which has consequences such as that the game may change stochastically to a loner game against Nature if one’s partner dies and that all payoffs tend to zero as the number of iterations tends to infinity. Results providing some insight into the evolution of cooperation are obtained when the number of iterations is large, including the conversion of cooperative dilemmas into situations in which cooperation is unambiguously favored.

Survival is an appropriate, direct measure of payoff (or utility) in evolution. Our analysis is similar to the analysis of diploid population-genetic models in that we have described interactions between individuals only in terms of fitness. What we have called strategies correspond to the alleles in a diploid population-genetic model. We have assumed that individuals’ strategies are fixed, meaning the same in every step of the game. For this sort of survival game, our results are general. They hold for any pattern of payoffs in the single-step matrix in Eq. (3). They can be applied to any biological, behavioral or ecological model for which the assumption of Markov survival is reasonable. They need not be pure strategies, for example. They could be mixed, as in Garay (2009), so long as they are the same in every step.

Our Markov assumption is twofold: (i) when an individual has a partner, its single-step survival probability depends on its strategy and on its partner’s strategy, and (ii) when an individual is alone, its single-step survival probability depends on its strategy. An individual’s multi-step survival probability is the product of its single-step survival probabilities given its situation in each step. The model of repeated predator attacks in Garay (2009) is precisely such a Markov survival model. In constructing the matrix in Eq. (3), we have further assumed (iii) that conditional on both of their strategies, which determine their individual survival probabilities, the two members of a pair survive each step independently of one another. If synergistic effects of partnership (Hauert et al., 2006; Kun et al., 2006) are defined as non-multiplicative, then assumption (iii) precludes synergy within a single step in our model. This would mean that all synergies in an *n*-step game result from the Markov process of survival for a given set of parameters (*a*, *b*, *c*, *d*, *a*_0_, *d*_0_).

We began with the motivation of Kropotkin (1902), whose work focused on vertebrates, but we note here that our model is not designed with any particular species in mind. None of our results rely on individuals possessing the intelligence or decision-making capability of vertebrates. It could be argued that many fundamental steps in the evolution of cooperation happened long before, perhaps many hundreds of millions of years before this kind of intelligence. The iterated survival game described in Section 2 may be applied in a variety of settings, but the evolutionary models of Sections 3 and 4 may be better suited to populations of single-celled organisms than complicated metazoans, because they assume haploid genetic transmission and individuals with fixed strategies.

Kropotkin inferred a connection between cooperation and survival under adverse conditions, and our work draws a distinction between games in which partnerships enhance survival and those in which having a partner is a liability. If there is a fitness advantage of having a partner, then less-cooperative behaviors which are advantageous in each single step of a game may become disadvantageous as the number of iterations increases. This can be true even if cooperative behaviors are strongly disfavored in the single-step game, as in the Prisoner’s Dilemma (e.g. Fig. 3A). But if partnerships do not significantly enhance survival, the opposite may occur. In this case, even cooperative behaviors which are favored in the single-step game can become disfavored as the number of iterations increases (e.g. Fig. 9A).

We close with a discussion of two examples which illustrate the effects of enhanced survival. We have identified two kinds of enhanced survival: for individuals and for pairs of individuals. These different boosts to survival are captured by the individual-survival increments of Eqs. (25) and (26), and by the presence of the pair-state eigenvalues of the Markov process in these same equations and their equivalents, Eqs. (23) and (24). Pairwise survival probabilities are crucial to understanding the structures of prolonged (large-*n*) games, which depend strongly the largest non-unit eigenvalue of the Markov survival process.

In the first example, having a partner of one particular strategy either increases or decreases the survival probability of the individual, but by an amount which does not depend on the strategy of the individual. We consider the two possibilities shown in Table 3, that either having an *A* partner increases survival or having a *B* partner decreases survival. In both cases, the change in survival (+*ϵ* or −*ϵ*) is the same for *A* individuals and *B* individuals. All other survival probabilities are equal to a baseline, *d*_0_. Thus *a* − *c* = 0 and *b* − *d* = 0. The single-step game is neutral. Figure 10 shows how *a*(*n*) − *c*(*n*) and *b*(*n*) − *d*(*n*) depend on *n* when *d*_0_ = 0.5 and *ϵ* = 0.1. In this case, *a*_0_ = *d*_0_ is the largest non-unit eigenvalue. In Fig. 10A, *a*(*n*) − *c*(*n*) > 0 for *n* ≥ 2 while *b*(*n*) − *d*(*n*) = 0 for all *n*. In Fig. 10B, *a*(*n*) − *c*(*n*) = 0 for all *n* while *b*(*n*) − *d*(*n*) < 0 for *n* > 2.

**Figure 10:**
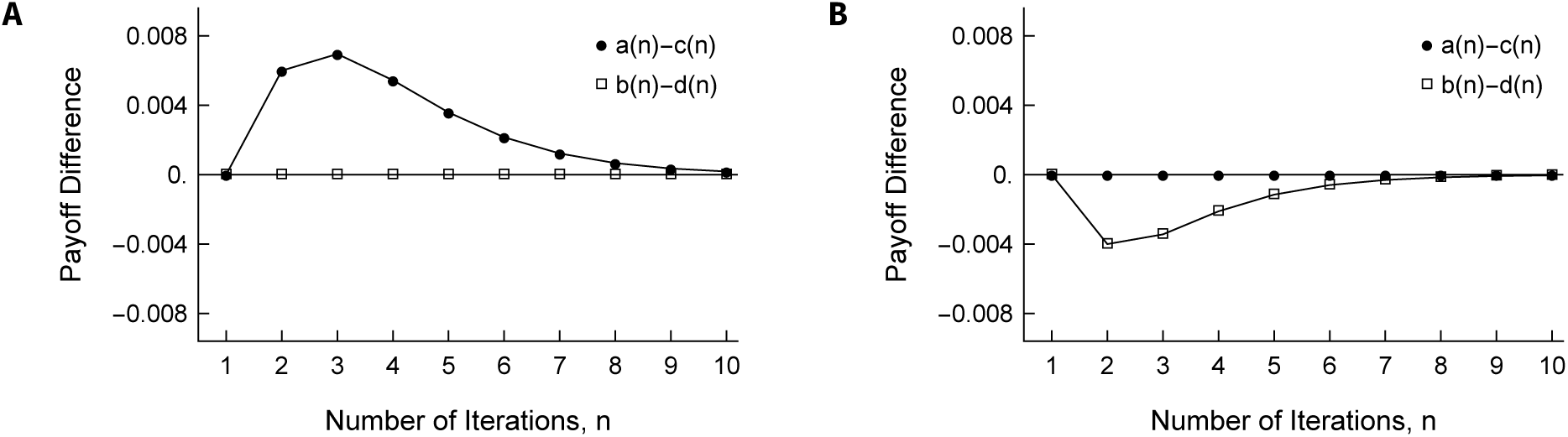
Values of the two key differences, *a*(*n*) − *c*(*n*) and *b*(*n*) − *d*(*n*), as a function of the number of iterations, for the two versions of the equal-payoff model depicted in Table 3, with baseline survival probability *d*_0_ = 0.5 and increment/decrement *ϵ* = 0.1. Panel A: *a* = 0.6, *b* = 0.5, *c* = 0.6, *d* = 0.5. Panel B: *a* = 0.5, *b* = 0.4, *c* = 0.5, *d* = 0.4. Both panels: *a*_0_ = *d*_0_ = 0.5.

**Table 3:** Two models in which individuals of type *A* and *B* have the same single-step probability of survival given the type of their partner, but in which the survival probability of any individual depends on the partner’s type. The format is as in Table 1 but with an additional column for the case in which an individual no longer has a partner. In the first model, having a partner of type *A* boosts the probability of survival of both types of individuals by an amount e above background (*d*_0_). In the second model, having a partner of type *B* lowers the probability of survival of both types of individuals by an amount *ϵ*.

| | $A$ | $B$ | No Partner |
| --- | --- | --- | --- |
| $A$ | $a = d_0 + \epsilon$ | $b = d_0$ | $a_0 = d_0$ |
| $B$ | $c = d_0 + \epsilon$ | $d = d_0$ | $d_0$ |

Table 3: Two models in which individuals of type $A$ and $B$ have the same single-step probability of survival given the type of their partner, but in which the survival probability of any individual depends on the partner's type.
| | $A$ | $B$ | No Partner |
| --- | --- | --- | --- |
| $A$ | $a = d_0$ | $b = d_0 - \epsilon$ | $a_0 = d_0$ |
| $B$ | $c = d_0$ | $d = d_0 - \epsilon$ | $d_0$ |

These results may be understood with reference to Eqs. (25) and (26), with *a*_0_ = *d*_0_. In the case of Fig. 10A, *a* − *a*_0_ = *c* − *d*_0_ = *ϵ* but, owing to the eigenvalues, the positive increment *a* − *a*_0_ is weighed more heavily than the negative increment *c* − *d*_0_ when *n* ≥ 2, so Eq. (25) gives *a*(*n*) − *c*(*n*) > 0 for *n* > 2. Equation (26) gives *b*(*n*) − *b*(*n*) = 0 because *b* − *a*_0_ = *d* − *d*_0_ = 0. Analogous arguments explain the case of Fig. 10B. In sum, the more-cooperative *A* is favored if it unilaterally increases the survival of its partner (Fig. 10A) and the less-cooperative *B* is favored if it unilaterally decreases the survival of its partner (Fig. 10B).

For the second example, we design an iterated survival game which expresses key ideas behind Model 2 of De Jaegher and Hoyer (2016) directly in terms of survival. Again, this is a model of a number of attacks on a pair of individuals. The individuals are identical in most respects. There is a public good which benefits all individuals equally, if it is preserved against attack. Each individual also has some private good which may benefit its partner too, but again only if it is preserved against attack. The difference is that cooperators (*A*) pay a cost and are immune to attacks whereas defectors (*B*) pay no cost and are susceptible to attacks. Thus, defectors may benefit when they are paired with cooperators. This is not a survival game because individuals do not die in the attacks. It is Markovian only in that each attack hits one member of the pair randomly with probability 1/2. Nonetheless, this behavioral and ecological model has roughly similar outcomes to those we have described here, when *n* corresponds to the number of attacks.

Consider a survival game in which the probability an individual survives each attack, or iteration, depends on its type and its partner’s type as follows.

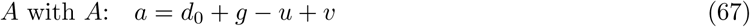

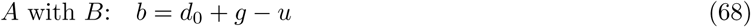

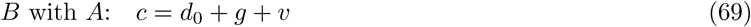

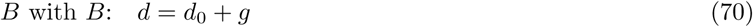

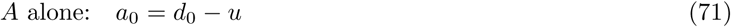

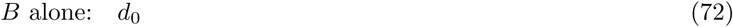

Here *u* is the cost of being a cooperator, v is the benefit an individual receives when its partner is a cooperator and *g* is the added benefit of simply having a partner. The parameters *d*_0_ and *g* might represent the private and public goods in De Jaegher and Hoyer (2016). The general versions of our model and Model 2 of De Jaegher and Hoyer (2016) each have seven free parameters, whereas the model above has five.

For this model *a* − *c* = *b* − *d* = −*u* < 0, so the cooperative type *A* is disfavored in the single-step game. In addition, *a*_0_ − *d*_0_ = −*u* < 0 which suggests that the cooperative type *A* might be disfavored for larger *n*. However, considering the terms in Eqs. (25) and (26) we have *a* − *a*_0_ = *c* − *d*_0_ = *g* + *v* > 0 and *b* − *a*_0_ = *d* − *d*_0_ = *g* > 0. If *a*^2^ is the largest non-unit eigenvalue then *a*(*n*) − *c*(*n*) will eventually become positive, and if *bc* is the next-largest eigenvalue then *b*(*n*) − *d*(*n*) will also eventually become positive.

A strong criterion for partnerships enhancing survival would be that *a*^2^, *bc* and *d*^2^ are all greater than *d*_0_. A weak criterion would be that only *a*^2^ > *d*_0_. If we adopt the strong criterion and further assume that *a*^2^, *bc* > *d*^2^, then the game would enhance survival if *d*^2^ > *d*_0_. This implies that

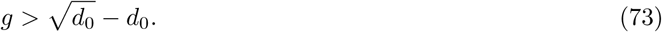

We have assumed throughout that *a*, *b*, *c* and *d* are all less than one, so it must also be true that

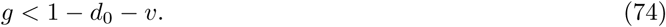

For the sake of illustration, set *g* equal to the midpoint between these two extremes, so that 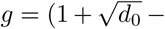 *v*)/2 − *d*_0_. Assume further that *v* > *u*, which means that the single-step game is a Prisoner’s Dilemma. Using the particular values *d*_0_ = 0.8, *u* = 0.02 and *v* = 0.03 gives the survival game shown in Table 4.

**Table 4:** Numerical example of the model defined in Eqs (67) through (72), assuming that *d*_0_ = 0.8, *u* = 0.02, *v* = 0.03 and with 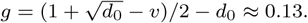

| | $A$ | $B$ | No Partner |
| --- | --- | --- | --- |
| $A$ | $a = 0.94$ | $b = 0.91$ | $a_0 = 0.78$ |
| $B$ | $c = 0.96$ | $d = 0.93$ | $d_0 = 0.80$ |

One would not immediately guess that the more cooperative type *A* would ever be favored in such a game. But as Fig. 11 illustrates, this does become true as the number of attacks increases. For *n* ≥ 32 both *a*(*n*) − *c*(*n*) and *b*(*n*) − *d*(*n*) are greater than zero, though clearly the prolonged game yields rather weak selection in this particular example. Note that we have not granted *A* any obvious advantage. As previously mentioned, cooperators are immune to attack in Model 2 of De Jaegher and Hoyer (2016). Here, instead, cooperation becomes favored as *n* increases due to the subtle advantages the more cooperative type *A* accrues because *AA* pairs survive best and *AB* second best among the three possible pairings, despite the direct disadvantage of *A* compared to *B* in *AB* pairs as well as in the loner state.

**Figure 11:**
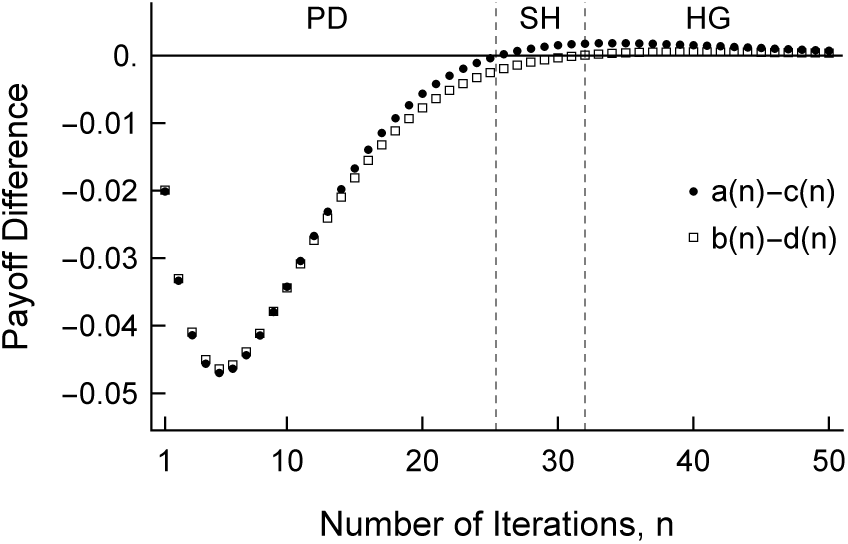
Values of the two key differences, *a*(*n*) − *c*(*n*) and *b*(*n*) − *d*(*n*) as a function of the number of iterations, for the game with payoffs shown in Table 4.

Rather than considering that individuals might change their strategies between steps of the game in response to their partners, we have focused on the evolution of hardwired, non-reactive strategies within populations. Among other results, we find the conditions under which cooperation will increase in frequency over time in both finite and infinite populations despite being disfavored in every step of the game. In the case of a finite population, this conclusion is based on a strong criterion for selective advantage, that the expected change in frequency of cooperation is positive over the entire range of possible frequencies.

## Acknowledgements

We thank Sabin Lessard and two anonymous reviewers for comments which greatly improved the work. We also thank Johannes Wirz for posing a question which led to the model presented in Table 3 and Fig. 10.

## Appendix

### Analysis of the Markov chain

Corresponding to the matrix in Eq. 3, we define the stochastic matrix

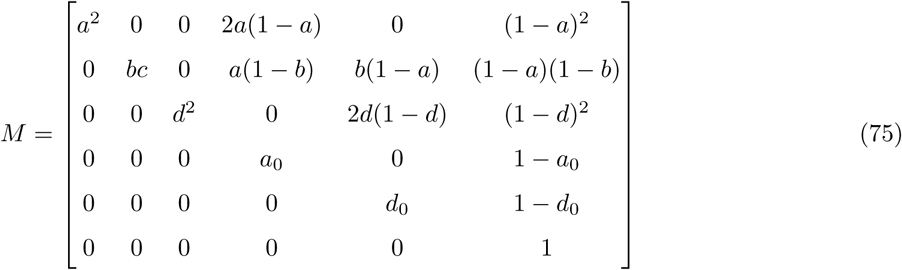

whose entries are the transition probabilities between the six possible states (*AA*, *AB*, *BB*, *A*, *B*, Ø). Thus, *M* describes a Markov process with a single absorbing state (Ø). When this is iterated *n* times, the probabilities of each of the six possible outcomes will be the entries of the *n*-step transition probability matrix *M^n^*. Because *M* is a triangular matrix, its eigenvalues are given by its diagonal elements. These are just the probabilities of remaining in each states over a single iteration. For reference, we reproduce Eq. (4)

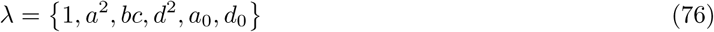

and further note that λ_2_, λ_3_ and λ_4_ are the probabilities that both individuals survive the iteration. Under the assumption that neither surviving nor perishing of individuals is guaranteed over a single iteration, which is to say 1 > *a*, *b*, *c*, *d*, *a*_0_, *d*_0_ > 0, λ_1_ = 1 is the largest eigenvalue and 1 > λ_2_, λ_3_, λ_4_, λ_5_, λ_6_ > 0.

From the standard theory of finite Markov chains (cf. section 2.12 in Ewens, 2004), if *r_i_* and *l_i_* are the corresponding right and left eigenvectors of *M*, normalized so that

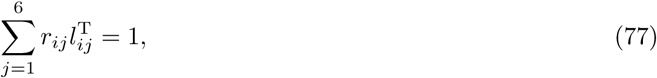

for each *i* ∈ (1, 2, 3, 4, 5, 6), then *M^*n*^* may be expressed in terms of its spectral decomposition

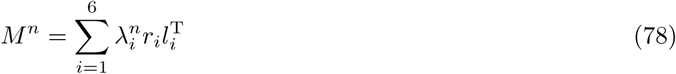

in which T denotes the transpose. With λ defined as above, the corresponding right and left eigenvectors of the matrix *M* are

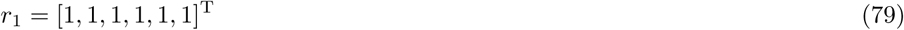

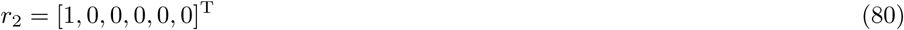

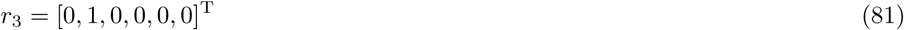

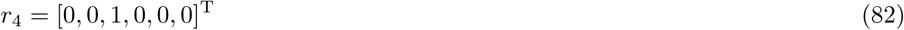

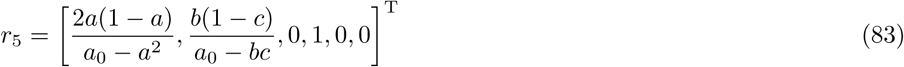

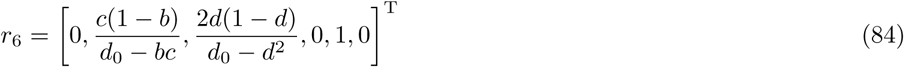

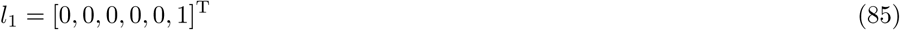

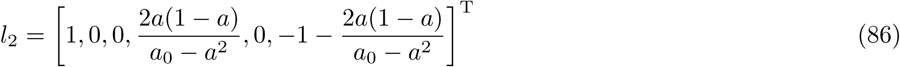

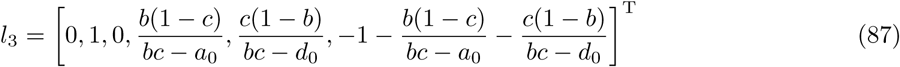

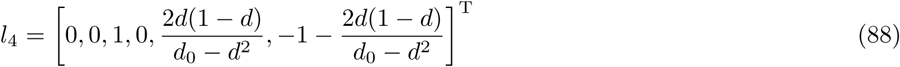

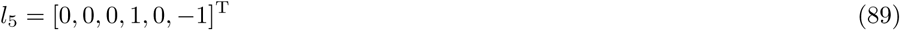

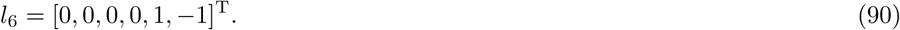

In terms of the notation here, the probabilities in the main text are

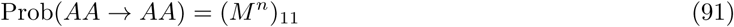

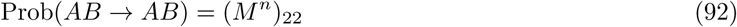

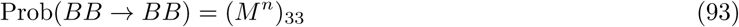

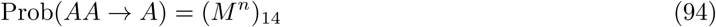

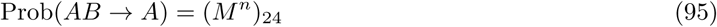

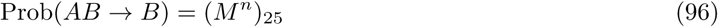

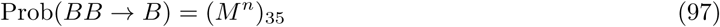

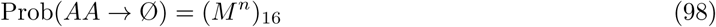

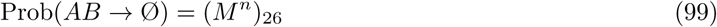

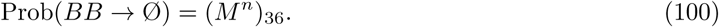

We can define the limiting matrix

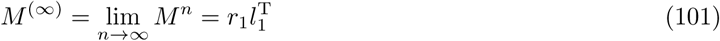

which has all six rows equal to [0, 0, 0, 0, 0, 1] and thus captures the fact that absorption in state Ø is guaranteed, if *n* is large enough, regardless of the starting state. Then we may write

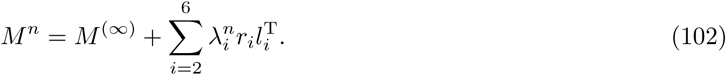

Because 1 > λ_*i*_ > 0 for *i* ∈ (2, 3, 4, 5, 6), the eventual rate of decay toward Mwill be depend on which λ_*i*_ is the leading non-unit eigenvalue of *M*. This, in turn, will depend on the values of *a*, *b*, *c*, *d*, *a*_0_ and *d*_0_. If *n* is very large, the sum in Eq. (102) will become dominated by a single term, namely the one involving the largest non-unit eigenvalue.

For example, if λ_2_ = *a*^2^ is the largest eigenvalue and *n* is very large,

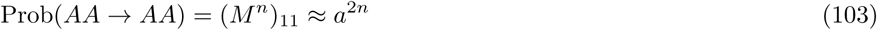

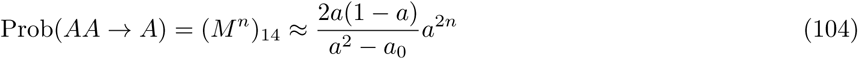

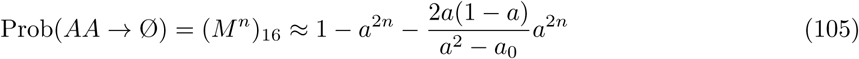

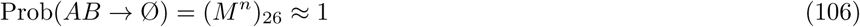

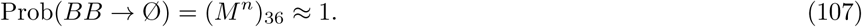

Then the effects of selection via the game reduce to

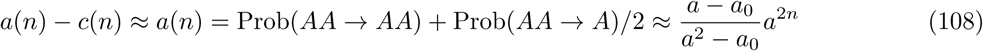

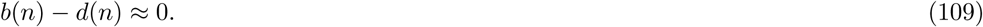

Our general results, Eqs. (23) and (24), are written to facilitate the direct inference of such conclusions.

### Derivation of the replicator equation

We assume a well-mixed population of infinite size. Well-mixed means specifically that initial partnerships in the game are formed at random in proportion to the current frequencies of *A* and *B*. The game is the only cause of death in the population and the only source of differences in fitness. Here we consider a continuous-time model. The replicator equation for the iterated survival game can be developed as follows. Changes in the frequencies (*x* and *y*) of *A* and *B* are given by a pair of differential equations

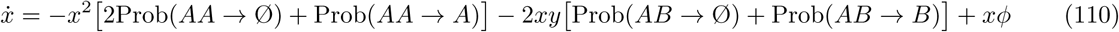

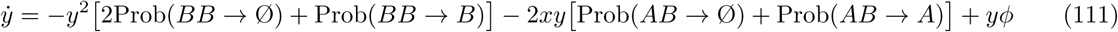

in which the factor of 2 inside the brackets in Eqs. (110) and (111) comes from the fact that both individuals die in the transition to state Ø. Individuals perish in the game, which causes *x* and *y* to decrease. This is balanced by reproduction, by the positive terms *xϕ* and *yϕ*. As there are only two types, *y* = 1 − *x*, so *x* + *y* = 1 and 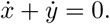 Then using these to solve for *ϕ* and substituting into Eq. (110) gives

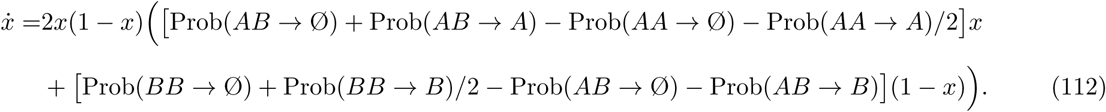

Note that pairs of terms in the brackets in Eq. (112) are related individual survival probabilities. For example, Prob(*AB* → Ø) + Prob(*AB* → *A*) = 1 − *c*(*n*). In addition, because the choice of time scale here is arbitrary, we may drop the leading factor of 2 in Eq. (112). Making these changes yields

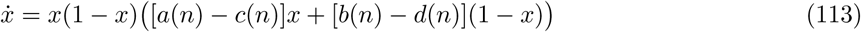

which is identical to Eq. (1) but now with payoffs from the *n*-step game.

### Classical population-genetic model

The replicator equation is a model of instantaneous, infinitesimal change in a population. Alternatively, we may follow the fates of pairs of individuals when generations are non-overlapping, as in classical models of diploid viability selection. The population is still assumed to be infinite but time is now measured in units of discrete generations. As a straightforward extension of classical models (Fisher, 1930; Haldane, 1932; Wright, 1931), we assume that the population well-mixed and that at the start of each generation it is composed of pairs *AA*, *AB* and *BB* in frequencies *x*^2^, 2*x*(1 − *x*) and (1 − *x*)^2^. The frequency of *A* is computed from the pair frequencies as

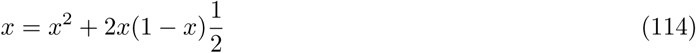

because both members of *AA* are *A* but only one member of *AB* is *A*. We emphasize that the accounting is of pairs. The total population is represented by

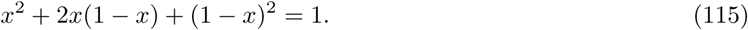

After selection, which is to say after the game, the population is composed of some intact pairs with types *AA*, *AB* or *BB* and some loners with types *A* or *B*. Their relative frequencies are

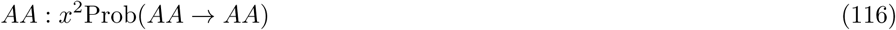

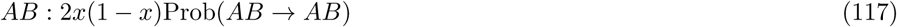

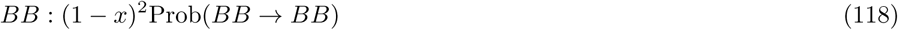

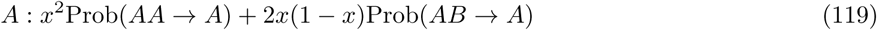

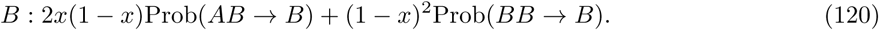

Loners must receive half the weight of intact pairs in this accounting. So, what remains of the total population is represented as

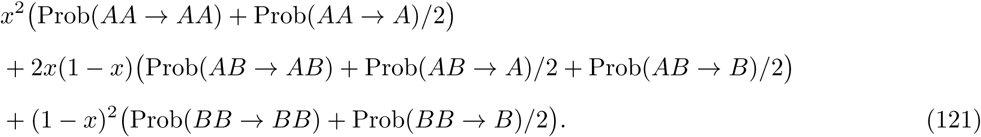

By definition, this is equivalent to the average fitness of the population. Using Eqs. (19) through (22), it may be written

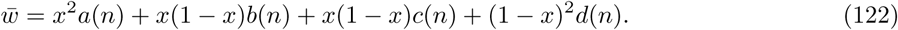

Note that is proportional to *ϕ* in Eqns. (110) and (111). Finally, accounting as in Eq. (114) for the fact that only one member of each intact *AB* pair is *A* and *w* in Eq. (122) for the average fitness, the frequency of *A* after the game is

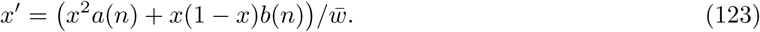

Then the change in the frequency of *A* over one generation, Δ*x* = *x*′ − *x*, is given by

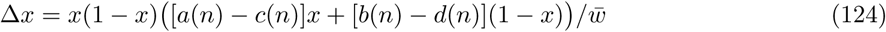

which is presented in the main text as Eq. (27).

